# Epigenetic control of nuclear mechanics and cellular migration via histone H3 lysine 9 methylation at Lamina-Associated Domains

**DOI:** 10.1101/2025.11.28.690983

**Authors:** Silvia Comunian, Valentin Petit, Guillaume Velasco, Vlada Zakharova, Cécile Jebane, Pauline Lahure, Véronique Joliot, Laurence Del Maestro, Magali Hennion, Guillaume Lamour, Jean-Baptiste Manneville, Emmanuèle Helfer, Ekaterina Boyarchuk, Slimane Ait-Si-Ali

**Author notes:** **Correspondence:** SA (;), EB (,). A51, Tour Montparnasse, 33 avenue du Maine 75015 Paris (Cécile Jebane).

## Abstract

Chromatin not only stores genetic information but also regulates nuclear mechanical properties. However, how epigenetic modifications, such as histone H3 lysine 9 trimethylation (H3K9me3), shape chromatin states at lamina-associated domains (LADs) and influence nuclear mechanics remains unclear. Here, we reveal an unexpected and paradoxical role of the pro-oncogenic H3K9 lysine methyltransferase (KMT) SETDB1, frequently overexpressed in many cancers, in LAD regulation. Rather than reinforcing heterochromatin, SETDB1 overexpression prevents SUV39H1-driven H3K9me3 accumulation at LADs, thereby disrupting the peripheral heterochromatin. Reducing SETDB1 levels, or disrupting its interaction with SUV39H1, enables SUV39H1 to restore LAD-localized H3K9me3 and re-establish an epithelial-like heterochromatin architecture. Heterochromatin reorganization upon SETDB1 reduction stiffens the nucleus and increases cell viscosity, thereby reducing cancer cell deformability and migratory capacity. Strikingly, these mechanical effects occur without major transcriptional changes, demonstrating that chromatin architecture itself critically shapes nuclear mechanics and cell motility. Our findings reveal a previously unrecognized role of the SETDB1–SUV39H1 axis in nuclear biomechanics, highlighting it as a potential determinant of cancer cell migration and metastatic processes.

## INTRODUCTION

The nucleus, the largest organelle in eukaryotic cells, is structurally defined by the nuclear envelope (NE), the underlying lamina, and chromatin (Stephens et al., 2019). Chromatin is spatially compartmentalized into transcriptionally active euchromatin (A compartment) and repressive, denser heterochromatin (B compartment) (Rowley & Corces, 2018). Heterochromatin is often enriched at the nuclear periphery within LADs, which are chromatin regions in direct molecular contact with the NE (van Steensel & Belmont, 2017). LADs are key regulators of chromatin organization and gene silencing, enriched in repressive histone marks on heterochromatin such as di- and tri-methylation of lysine 9 on histone H3 (H3K9me2 and H3K9me3, respectively) (Bian et al., 2013; Briand & Collas, 2020; Guelen et al., 2008).

Post-translational modifications (PTMs) of histones are essential for chromatin architecture. H3K9 methylation is a repressive mark catalyzed by specific lysine methyltransferases (KMTs), notably members of the SUV39 subfamily (Mozzetta et al., 2015). G9A and GLP primarily catalyze mono-and di-methylation, whereas SETDB1 and SUV39H1 mediate H3K9 tri-methylation (Mozzetta et al., 2015). SETDB1 can catalyze all three methylation states in both euchromatin and heterochromatin (Mozzetta et al., 2015), while SUV39H1 catalyzes di- and tri-methylation, with preferential enrichment in constitutive heterochromatin. These KMTs often act cooperatively; for instance, SETDB1 interacts with G9A, GLP, and SUV39H1 to promote transcriptional repression at transposable elements such as intracisternal A particle (IAPs) and long interspersed elements (LINEs) (Bulut-Karslioglu et al., 2014; Fritsch et al., 2010; Karimi et al., 2011; Liu et al., 2014; Padeken et al., 2022). This cooperation extends to the establishment and spreading of H3K9me3 at telomeric and pericentromeric regions, and at LADs (Gauchier et al., 2019; Loyola et al., 2009; Towbin et al., 2012). However, the specific roles and interactions of these enzymes at LADs remain insufficiently understood.

Dysregulation of KMTs and their associated epigenetic marks is frequently linked to pathological conditions, including cancers (Mozzetta et al., 2015). SETDB1 is overexpressed in several cancers, notably non-small cell lung cancer (NSCLC), where it is associated with increased metastasis and poor prognosis (Cruz-Tapias et al., 2019). In NSCLC, and in many epithelial cancer types (Strepkos et al., 2021), SETDB1 represses epithelial genetic programs, including genes governing cell-cell adhesion and cell migration (Zakharova et al., 2022). In normal human bronchial epithelial cells (NHBE), H3K9me3 is enriched at LADs, contributing to nuclear structure and transcriptional gene silencing (Keenan et al., 2024; Zakharova et al., 2022). This heterochromatin organization is disrupted in NSCLC (Zakharova et al., 2022), suggesting that SETDB1 overexpression may lead to H3K9me3 redistribution and altered nuclear structure.

In addition to regulating transcription, chromatin organization, including the spatial distribution of H3K9me3, plays a key role in determining nuclear mechanical properties (Nava et al., 2020). During metastasis, cancer cells undergo the epithelial-to-mesenchymal transition (EMT), which enables them to migrate through dense physical environments such as stroma, blood vessel walls, and ultimately the tissue at the secondary site (Chaffer & Weinberg, 2011; Wirtz et al., 2011). This migration requires significant nuclear deformation as the nucleus is the largest and stiffest cellular organelle (Gensbittel et al., 2021; Golloshi et al., 2022; Harada et al., 2014; Nia et al., 2020). Therefore, nuclear stiffness has emerged as a critical determinant of metastatic potential. Given that H3K9me3 enrichment at LADs influences chromatin compaction (Feng et al., 2020), we hypothesized that it may regulate nuclear mechanics and, consequently, cancer cell motility and migration.

Our previous work showed that a complete SETDB1 knockout disrupts cell migration through combined effects on EMT gene expression and chromatin reorganization at H3K9me3-marked LADs (Zakharova et al., 2022). The current study demonstrates a new mechanism: reducing SETDB1 levels is sufficient to remodel chromatin architecture and thereby alter the biophysical properties of cells. Partial knockdown of SETDB1 has little impact on gene expression, yet it still compromises migration through chromatin-based changes in nuclear mechanics. Specifically, restoration of H3K9me3 at LADs leads to increased nuclear stiffness and circularity, and cellular viscosity, accompanied by enlarged nuclear and cellular sizes.

In this study, we demonstrate that the interplay between SETDB1 and SUV39H1 regulates the distribution H3K9me3 at LADs in NSCLC cells. Based on this interplay, we propose a model in which overexpressed SETDB1 limits SUV39H1-dependent increase of H3K9me3 at LADs. Conversely, SETDB1 depletion or mutation releases SUV39H1, enabling the restoration of H3K9me3 at the nuclear periphery. Overexpression of SUV39H1 in NSCLC cells recapitulates SETDB1 loss, reinforcing the cooperative mechanism.

In summary, our findings highlight a non-canonical role for SETDB1 in epigenetically regulating nuclear stiffness via its interference with SUV39H1 at LADs. By modulating the spatial distribution of H3K9me3, SETDB1 influences the mechanical properties and invasive behavior of lung cancer cells. By interfering with SUV39H1 function, SETDB1 exhibits a novel, most likely non-transcriptional role in shaping nuclear biomechanics. These findings support the hypothesis that targeting the SETDB1–SUV39H1 axis could offer a novel therapeutic strategy to limit cancer cell migration and metastasis.

## RESULTS

### SETDB1 reduction impairs migration in SETDB1-overexpressing cancer cells with minimal impact on gene expression

SETDB1 is overexpressed in the non-small cell lung cancer (NSCLC) cell line A549 (Ctrl) compared to normal NHBE lung epithelial cells (Guo et al., 2014; Weirich et al., 2021; Zakharova et al., 2022). Our previous study showed that a stable SETDB1 loss-of-function induced by a knockout (KO) approach in A549 cells triggers extensive transcriptional reprogramming, including de-repression of epithelial gene programs, and markedly impairs cell migration (Zakharova et al., 2022). However, given the widespread transcriptional effects of SETDB1 KO, it remains unclear whether the observed cell migration defects arise directly from alterations in H3K9me3 deposition or indirectly from secondary transcriptional changes. To address this uncertainty, we employed an shRNA-mediated knockdown (KD) approach to partially reduce SETDB1 levels. This strategy allowed us to test whether a partial reduction of SETDB1 is sufficient to impair cell migration while minimizing transcriptional perturbations.

To this end, we developed a doxycycline (dox)-inducible shRNA system targeting SETDB1 (SETDB1 KD) and examined cells either untreated (0) or under 3-day (3) and 7-day (7) dox treatment conditions (**Fig. 1A**). As an additional control, we used a non-targeting shRNA (shCTR) to exclude shRNA off-target effects. The SETDB1 KD system reduced SETDB1 mRNA levels by approximately 80% (**Fig. 1B**) and protein levels by ∼50% after 7 days of dox treatment, while the control shCTR cell line showed no changes in SETDB1 protein and mRNA levels upon dox addition (**Fig. S1 A, B** and **D**).

**Figure 1.**
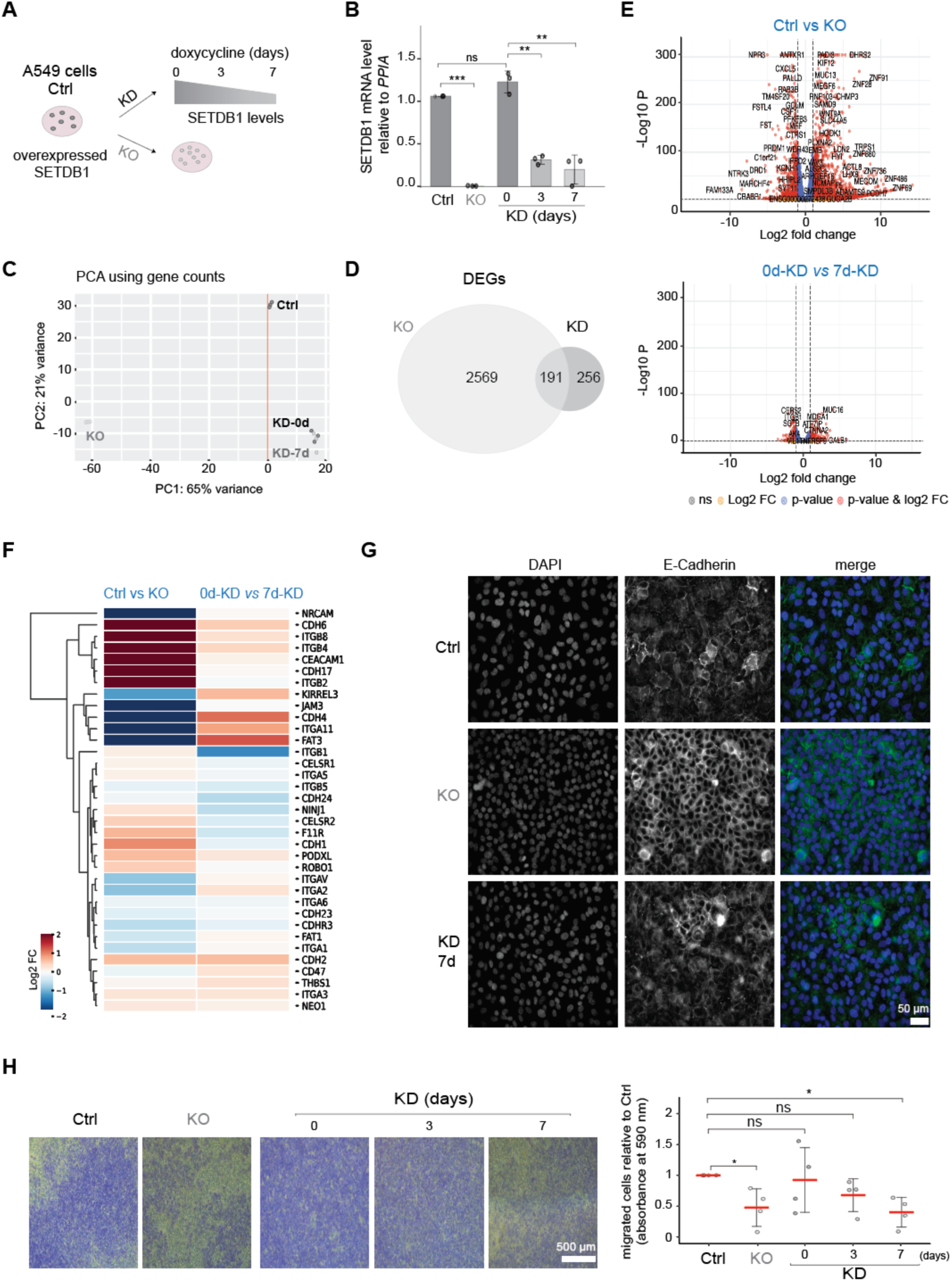
Reduction of SETDB1 levels impairs cell migration with minimal impact on global gene expression. **A.** Schematic representation of the cell lines exhibiting different SETDB1 expression levels. A549 Ctrl cells exhibit endogenous overexpression of SETDB1. SETDB1 KO cells show complete loss of SETDB1(Zakharova et al., 2022). In the inducible shRNA-mediated knockdown model (SETDB1 KD), cells are either untreated (0) or treated with doxycycline for 3 or 7 days (3d, 7d), resulting in a progressive reduction of SETDB1 levels. **B.** SETDB1 mRNA levels measured by RT-qPCR and normalized to *PPIA*. Values are shown relative to A549 cells (Ctrl). Bars represent the mean of N=3 biological replicates +/− SD error bars., with individual data points indicated. Statistical significance was assessed using an unpaired t-test; ns indicates p > 0,05, * indicates p < 0.05, ** indicates p < 0.01, *** indicates p < 0.001. **C.** Principal Component Analysis (PCA) based on gene expression profiles for A549 cells (Ctrl), SETDB1 KO, SETDB1 KD 0 (no doxycycline), and SETDB1 KD 7d-dox. **D.** Venn diagram of differentially expressed genes (DEGs). Comparison of Control vs SETDB1 KO (light grey) and SETDB1 KD 0 vs 7d-dox (dark grey). DEGs were selected based on a specified log2 fold change (log2FC) threshold: log2FC > +1 or <-1. **E.** Volcano plots showing -log10(p-value) (-Log10 P) versus log2 fold change for gene expression comparisons: Control vs SETDB1 KO and SETDB1 KD 0 vs 7d-dox. **F.** Heatmap of log2 fold changes for genes associated with cell migration, comparing Ctrl vs SETDB1 KO and SETDB1 KD 0 vs 7d-dox. Migration-related genes were selected from Gene Ontology database (<u>GO:0016477</u>). **G.** Representative immunofluorescence images of Ctrl, SETDB1 KO, SETDB1 KD 7 days-dox cells stained with anti-CDH1 (E-cadherin, green). Nuclei are counterstained with DAPI (blue). Merged images show overlay of CDH1 and DAPI signals. Scale bar: 50 µm. **H.** *Left:* Representative images from Transwell migration assays for Control, SETDB1 KO, and SETDB1 KD (0, 3, 7 days-dox). Migrated cells are stained in violet; images acquired at 40× magnification. Scale bar: 500 µm. *Right:* Quantification of migrated cells by absorbance measurement at 590 nm. Data represents N=4 replicates; red lines indicate the sample mean +/−SD error bars. Statistical analysis performed via unpaired t-test; ns indicates p > 0,05, * indicates p < 0.05, ** indicates p < 0.01, *** indicates p < 0.001.

Next, we evaluated how SETDB1 reduction affects gene expression by RNA sequencing and comparing (i) Control vs. SETDB1 KO cells and, (ii) untreated SETDB1 KD (0) vs. 7d dox-treated cells. Principal component analysis (PCA) revealed that SETDB1 KO cells were distinctly separated from all other conditions, whereas the transcriptome of SETDB1 KD remained closely aligned with the control group (**Fig. 1C**). Although SETDB1 mRNA was effectively reduced in the KD model, expression of known SETDB1 target genes such as *ATF7IP* and *PCDHB6* remained mostly unaffected, in contrast to their strong deregulation in SETDB1 KO cells (**Fig. S1D**).

Differential gene expression analysis further confirmed a significantly lower transcriptional shift in KD cells compared to KO cells, which had a higher number of differentially expressed genes (DEGs) (**Table S1**; **Fig. 1D**). The DEGs observed in the 7d dox-treated cells compared to untreated control showed lower log_2_ fold changes and less robust statistical significance compared to changes in KO condition (**Fig. 1E**). The changes in SETDB1 KD were comparable to those in the control shCTR (**Fig. S1C**). When focusing on genes associated with cell migration (GO:0016477), SETDB1 KO induced a marked transcriptional response, whereas KD resulted in only mild changes (**Fig. 1F**). Gene Set Enrichment Analysis (GSEA) confirmed that 50 % SETDB1 KD had different effects on gene expression than total SETDB1 KO. While some pathways remained unaffected by partial SETDB1 reduction (*e.g.,* the MYC-targets, **Fig. S1E**), others shifted in response to shRNA-mediated KD. Interestingly, several pathways exhibited opposing regulation between the SETDB1 KO and KD conditions. A striking example is the WNT/β-catenin signaling pathway, which was oppositely modulated in the two systems (**Fig. S1E**). For cell migration-related pathways, the normalized enrichment score (NES) for the apical junction gene set was increased in both conditions but reached statistical significance only in KO cells, where cell-cell adhesion is more impacted (**Fig. S1F**). This was corroborated by immunofluorescence showing stronger E-cadherin (CDH1) staining in KO cells, while both control and KD cells displayed weaker junctional signals (**Fig 1G, S1G**). Similarly, the immunostaining of Paxillin, a focal adhesion marker, revealed altered adhesion in KO cells but not in KD cells, highlighting distinct cellular consequences between the two models (**Fig. S1 H-I**).

Intriguingly, despite a partial reduction of SETDB1 levels and a mild effect on migration-related genes, SETDB1 KD (7d) cells exhibited impaired invasive capacity in Transwell migration assays, similar to SETDB1 KO cells (**Fig. 1H**). Additionally, both SETDB1 KO and SETDB1 KD cells showed reduced migratory ability in wound healing assays (**Fig. S1J**). Notably, proliferation defects were detected only in KO and not in SETDB1 KD cells (**Fig. S1K)**, indicating that the delayed wound closure in SETDB1 KD cells was not due to impaired proliferation.

These results demonstrate that partial reduction of SETDB1 levels is sufficient to compromise cell motility. We therefore concluded that SETDB1 is a key regulator of migration of A549 NSCLC cells, and that its function in motility is not strictly dependent on its role in transcriptional regulation. While a total SETDB1 KO triggers broad gene expression changes and disrupts cell-cell adhesions and focal adhesions, a partial KD is sufficient to impair migration without inducing major transcriptional shifts. This uncoupling of migratory defects from transcriptional deregulation suggests that SETDB1 may modulate cancer cell motility through non-transcriptional mechanisms, potentially involving chromatin organization and/or nuclear mechanics.

### SETDB1 loss of function affects H3K9me3 distribution at nuclear periphery without affecting LADs identity

To explore the chromatin organization effects of SETDB1 partial depletion by KD, we focused on the global distribution of H3K9me3, which we measured using high-throughput microscopy. We found that while global H3K9me3 levels were markedly reduced in SETDB1 KO cells, consistent with our previously published data (Zakharova et al., 2022), they remained largely unchanged upon SETDB1 KD (**Fig. S2A**). Unexpectedly, however, we detected a redistribution of H3K9me3 toward the nuclear periphery (**Fig. 2A**), as reflected by an increased peripheral-to-total H3K9me3 ratio (**Fig. 2B** and **Fig. S2 B-C**). This suggests that reducing SETDB1 levels is sufficient to promote H3K9me3 enrichment at LADs.

**Figure 2.**
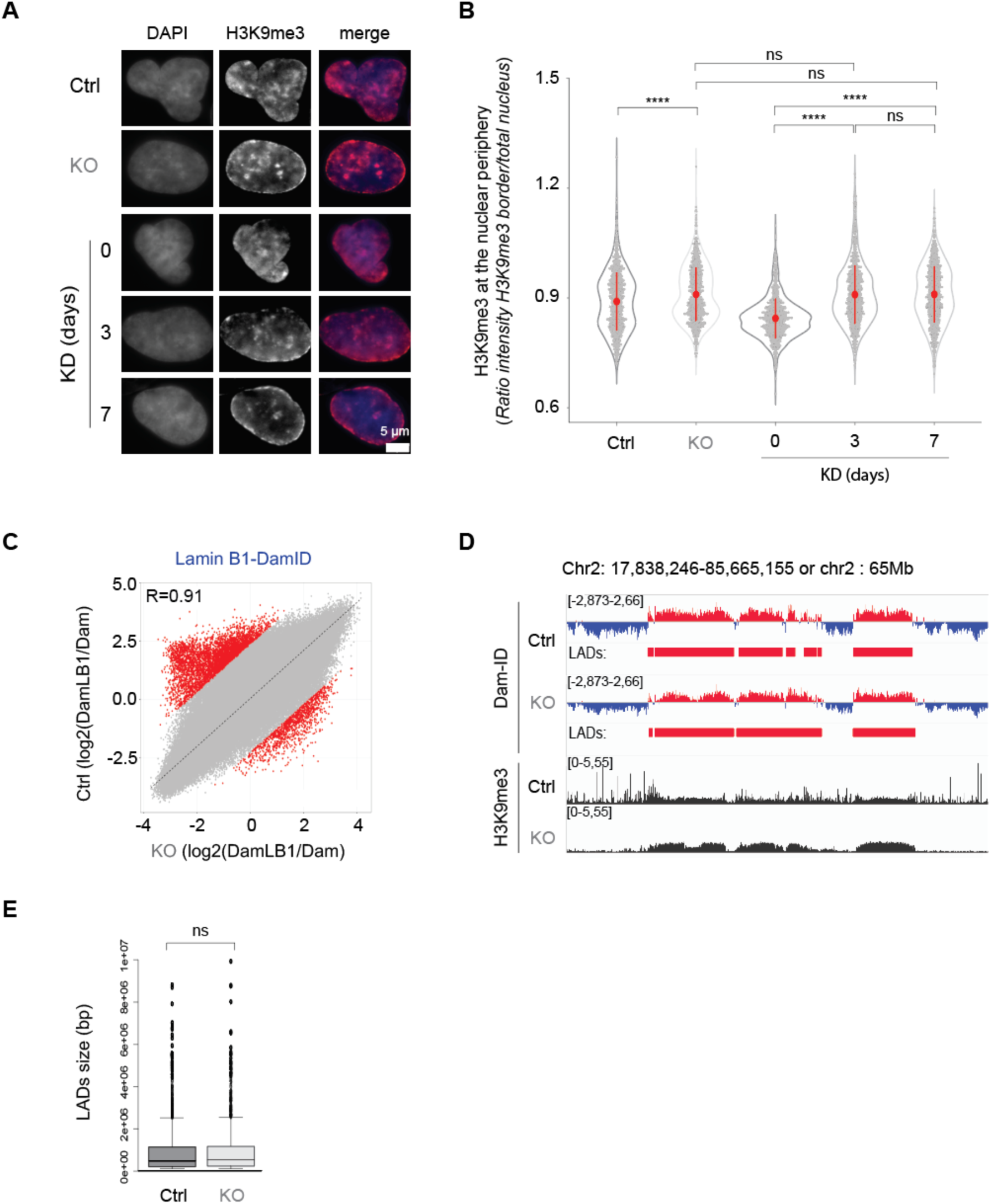
SETDB1 regulate the H3K9me3 distribution at the nuclear periphery but does not impact the LADs identity. **A.** Representative immunofluorescence images of A549 cells (Ctrl), SETDB1 KO, SETDB1 KD 0, 3d and 7d dox-treated cells stained with anti-H3K9me3 antibody. DNA was labelled with DAPI. In the merge the DAPI is represented in blue and the H3K9me3 in red. Scale bar: 5 µm. **B.** Violin plot showing the quantification of H3K9me3 signal at the nuclear periphery measured with the automated microscope (Operetta) corresponding to the phenotype observed in panel A. Each point represents the ratio between the intensity of the signal at the nuclear periphery (defined by DAPI) and the total nuclear H3K9me3 signal intensity. 1000 cells were randomly picked over more than 10 000 cells per condition. The red dot indicates the mean of N=3 biological replicates +/− SD error bars. Statistical significance was determined using the Wilcoxon test; ns indicates p > 0,05, * indicates p < 0.05, ** indicates p < 0.01, *** indicates p < 0.001, **** indicates p < 0.0001. **C.** Scatter plots showing the correlation between normalized read counts (log_₂_[Dam-LB1/Dam]) from DamID-seq experiments performed in Ctrl and SETDB1 KO cells, using a bin size of 10 kb. The Spearman correlation coefficient (R) is indicated on the plot. The dashed line represents the fitted correlation line. Outliers exceeding ±4 standard deviations from the correlation line are highlighted in red and denote genomic regions specifically enriched in either Ctrl or SETDB1 KO cells. **D.** Integrated Genome Viewer (IGV) tracks displaying DamID-seq and ChIP-seq data from Ctrl and SETDB1 KO cells. Tracks represent *Dam-*Lamin B1 (DamID) and H3K9me3 (ChIP_H3K9me3) normalized signals, respectively. Genomic regions corresponding to lamina-associated domains (LADs) are highlighted in red, while inter-LAD regions are shown in blue. Although LADs are broadly conserved across the depicted region of chromosome 2 (Chr2), the figure reveals a pronounced increase in H3K9me3 signal at LADs in SETDB1 KO cells compared to control cells. H3K9me3 ChIP-seq data were obtained from reference [18]. **E.** Boxplot comparing the size of LADs, measured in base pairs, between Ctrl and SETDB1 KO cells. Statistical analysis indicates no significant difference (ns) between the two conditions.

To assess whether the redistribution of H3K9me3 in absence of SETDB1 is accompanied by changes in LAD genomic content, defined as large (0.1-10 Mb) genomic regions in contact with the nuclear lamina (NL), we performed DamID-seq using a Dam-Lamin B1 fusion protein to map LADs. Despite the pronounced H3K9me3 redistribution in SETDB1 KO cells, the global LAD landscape remained largely unchanged, with a comparable number of LADs identified in Control and KO conditions (1,527 vs. 1,461, respectively). Most LADs overlapped with previously defined H3K9me3-enriched domains (**Fig. 2 C-D**), and their sizes were similar between conditions (**Fig. 2E**). However, 47 and 51 large genomic regions (≥100 kb; FC > 1.5, p-adjust < 0.05) lost or gained NL association, respectively, in SETDB1 KO cells, encompassing 866 and 274 genes (GENCODE v47; **Fig. S2 D-E**). These LAD changes were limited across individual chromosomes, with chromosome 19 exhibiting the most notable reduction in LAD content, coinciding with regions under transcriptional regulation by SETDB1 (Zakharova et al., 2022) (**Fig. S2 F-G**).

### The SETDB1-SUV39H1 balance controls H3K9me3 distribution at nuclear periphery

Our previous study (Zakharova et al., 2022) showed that the KD of SUV39H in SETDB1 KO cells abolished the H3K9me3 enrichment at several LADs, as measured by ChIP-qPCR, suggesting that SUV39H is involved in establishing H3K9me3 enrichment at LADs. Consistently, siRNA-mediated KD of SUV39H in both control and SETDB1 KO cells (**Fig. S3 A-B**) significantly reduced H3K9me3 enrichment at the nuclear periphery in SETDB1 KO cells and further decreased H3K9me3 levels at the nuclear periphery in A549 control cells (**Fig. S3 C-D**).

To further test whether the balance between SETDB1 and SUV39H levels is important for establishing H3K9me3 enrichment at LADs, we overexpressed either wild-type (WT) SUV39H1 or a catalytic mutant of SUV39H1 in A549 Ctrl cells. Overexpression of WT SUV39H1 alone was sufficient to increase H3K9me3 levels at the nuclear periphery, whereas the SUV39H1 mutant had no effect (**Fig. 3 A-B**). Notably, WT SUV39H1 overexpression induced a slight increase in total H3K9me3 levels, but not that of the catalytically-inactive form (**Fig. S3E**). Importantly, this increase in H3K9me3 levels at the nuclear periphery induced by SUV39H1 overexpression mimicked the redistribution observed upon SETDB1 depletion (**Fig. 2A** and **Fig. 3A**).

**Figure 3.**
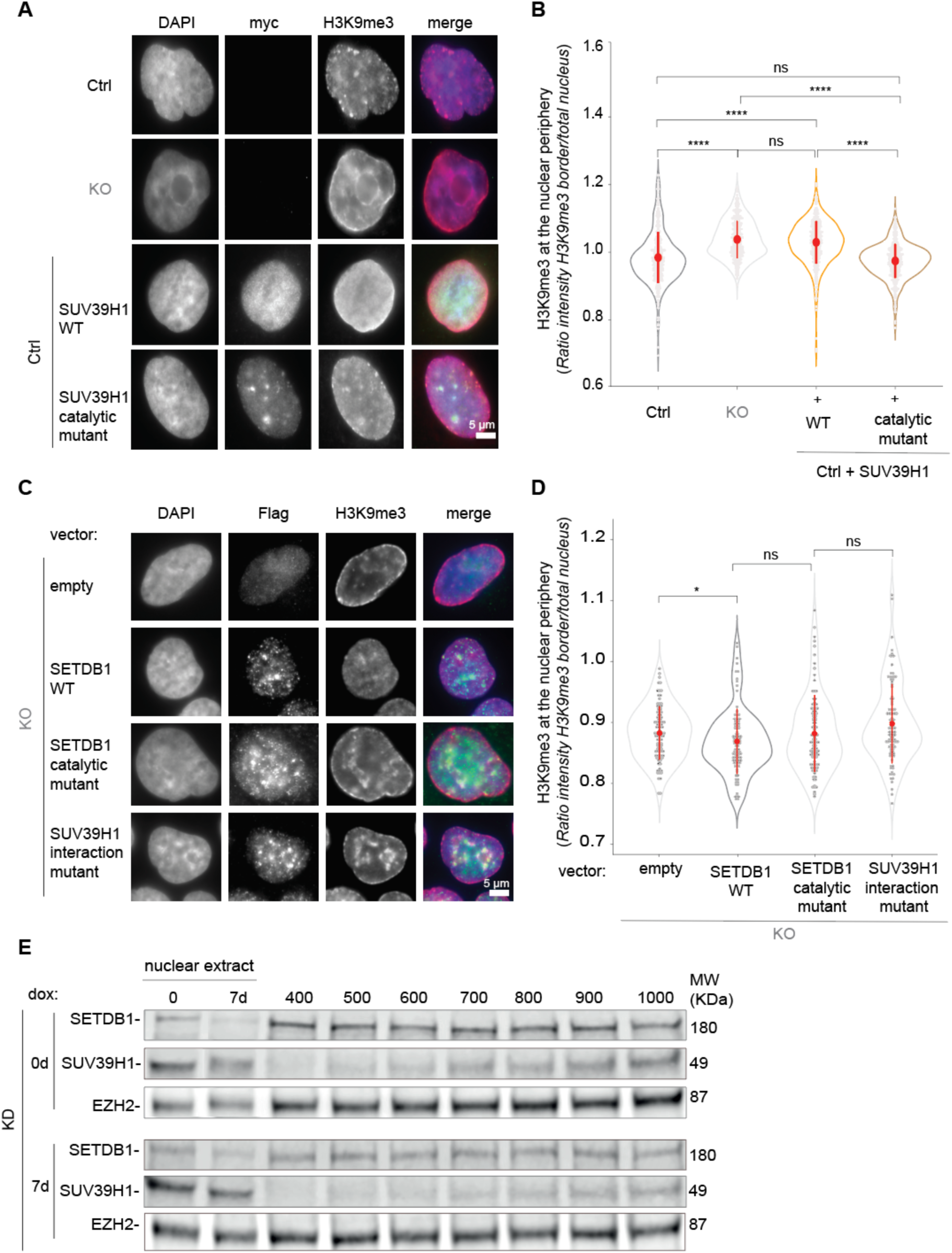
The SETDB1-SUV39H1 balance controls H3K9me3 distribution at the nuclear periphery. **A.** Representative immunofluorescence images of Ctrl, SETDB1 KO, Ctrl + SUV39H1 WT, Ctrl + SUV39H1 catalytic mutant. Cells were stained with anti-H3K9me3 antibody and positive cells expressing exogenous SUV39H1 WT or catalytic mutant constructs were stained with anti-MYC antibody. The nuclei were labelled with DAPI. The merge images show the DAPI in blue, MYC in green and H3K9me3 in red. Scale bar: 5µm. **B.** Violin plot showing the quantification of H3K9me3 signal at the nuclear periphery measured with the automated microscope (Operetta) corresponding to the phenotype observed in panel A. Cells analyzed are considered positive based on the value of the total intensity of the MYC signal inside the nucleus. Each point represents the ratio between the intensity of the signal at the nuclear periphery (defined by DAPI) and the total nuclear H3K9me3 signal intensity. 1000 cells were randomly picked over more than 10 000 cells per condition. The red dot indicates the mean of N=3 biological replicates +/− SD error bars. Statistical significance was determined using the Wilcoxon test; ns indicates p > 0,05, * indicates p < 0.05, ** indicates p < 0.01, *** indicates p < 0.001, **** indicates p < 0.0001. **C.** Representative immunofluorescence images of SETDB1 KO + empty vector, + SETDB1 WT, + SETDB1 catalytic mutant or SUV39H1 interaction mutants. Cells were stained with anti-H3K9me3 and anti-FLAG antibodies. Flag is used to identify the positive cells. DNA was labelled with DAPI. The merged images show DAPI in blue, H3K9me3 in red and the FLAG in green. Scale bar: 5 µm. **D.** Violin plot showing the quantification of H3K9me3 signal at nuclear periphery measured with the automated microscope (Operetta) corresponding to the phenotype observed in panel C. Cells analyzed are considered positive based on the value of the total intensity of the FLAG signal within the nucleus. Each point represents the ratio between the intensity of the signal at the nuclear periphery (defined by DAPI) and the total nuclear H3K9me3 signal intensity. 100 cells were randomly picked over more than 3000 cells per condition. The red dot indicates the mean of N=3 biological replicates +/− SD error bars. Statistical significance was determined using the Wilcoxon test; ns indicates p > 0,05, * indicates p < 0.05, ** indicates p < 0.01, *** indicates p < 0.001, **** indicates p < 0.0001. **E.** Solubility test for SUV39H1 in SETDB1 KD 0 days-dox (top) and SETDB1 KD 7 days-dox (bottom) cells using increasing NaCl concentration (400 to 1000 mM). SETDB1 is controlled to show the efficiency of the dox treatment. EZH2 is used as control to show that there is no change in solubility between 0 and 7 days-dox SETDB1 KD. On the left the total nuclei extract shows that the level of SUV39H1 in 0 and 7 days-dox SETDB1 KD cells.

Taken together, these results suggest that SUV39H1 is the key KMT responsible for H3K9me3 accumulation at LADs, particularly when SETDB1 is depleted in A549 cells, and that the balance between SETDB1 and SUV39H1 levels is crucial for establishment and/or maintenance of H3K9me3 enrichment at LADs.

To elucidate the mechanism by which the interplay between SETDB1 and SUV39H1 regulates H3K9me3 enrichment at LADs, we first assessed whether the catalytic activity of SETDB1 is required by performing rescue experiments in SETDB1 KO cells. Re-expression of WT SETDB1 in these cells restored a uniform nuclear H3K9me3 distribution, comparable to that of control A549 cells (**Fig. 3 C-D**, top two panels). In contrast, re-expression of an inactive SETDB1 catalytic mutant failed to reverse the KO phenotype (**Fig. 3C**, third row and **Fig. 3D**). These findings indicate that the enzymatic activity of SETDB1 is essential for the loss of H3K9me3 enrichment at LADs in A549 cells, without significantly affecting the total level of H3K9me3 (**Fig. S3F**).

SETDB1 and SUV39H1 are known to interact both physically and functionally (Fritsch et al., 2010). Since the balance between the protein levels of these two KMTs is instrumental in establishing H3K9me3 enrichment at LADs, we hypothesized that their interaction might be important. Recently, we showed that SETDB1 undergoes auto-methylation at two lysines within histone-mimic motifs, and these two methylatable lysines are necessary for its efficient interaction with several chromodomain-containing proteins, including SUV39H1 (Cruz-Tapias, 2025). Notably, these two lysines are conserved and methylated in human SETDB1 (K1186 and K1194) (Cruz-Tapias, 2025; Guo et al., 2014). We therefore performed rescue experiments using a lysine-to-alanine (K to A) non-methylatable mutant (K1186A and K1194A, interaction mutant, **Fig. S3G**) that shows reduced interaction with SUV39H1 (Cruz-Tapias, 2025). Notably, it was shown that the substitution of these two lysines does not affect the catalytic activity of SETDB1, but impacts its genome-wide distribution (Cruz-Tapias, 2025). Importantly, this SETDB1-SUV39H1 interaction mutant was not able to restore H3K9me3 distribution in SETDB1 KO cells (**Fig. 3C**, fourth row), in contrast to WT SETDB1, and similarly to SETDB1 catalytically inactive mutant (**Fig. 3 C-D**). Altogether, these data strongly suggest that the interplay between SETDB1 and SUV39H1, in cancer cells overexpressing SETDB1, limits SUV39H1-mediated establishment of H3K9me3 enrichment at LADs.

We next hypothesized that in the absence of its interaction with SETDB1, SUV39H1 might change its interacting partners, resulting in changes in its biochemical behavior, for example, chromatin-binding properties. We therefore performed a chromatin solubility assay for SUV39H1 in A549 control, SETDB1 KD, and SETDB1 KO cells using increasing salt concentrations. We found that SUV39H1 was extracted with a lower salt concentration, and therefore less tightly bound to the chromatin, in control cells where SETDB1 is overexpressed, compared to SETDB1 KD 7d-dox and KO cells (**Fig. 3E** and **Fig. S3H**).

Based on these findings, we suggest that SETDB1 overexpression in cancer cells shifts SUV39H1 into a more soluble compartment, away from LADs, thereby limiting SUV39H1 activity at LADs and possibly in other heterochromatin compartments. In SETDB1 KO cells and SETDB1 KD conditions, this retention is lifted, allowing SUV39H1 to access LADs, leading to increased H3K9me3 levels at LADs. These results suggest that SETDB1 overexpression plays a critical role in regulating SUV39H1 activity and preventing H3K9me3 enrichment at LADs, which may confer a selective advantage to cancer cells. This advantage could include changes in nuclear mechanical properties, that might affect cell migration potential in confined environments.

### H3K9me3 levels at LADs impact cellular viscosity and nuclear rigidity

Drawing from our findings that the SETDB1-SUV39H1 axis modulates cancer cell motility also through non-transcriptional mechanisms (**Fig. 1** and **Fig. S1**), we propose that alterations in H3K9me3 levels at LADs may affect the biophysical properties of cells. To investigate this hypothesis, we observed the entry of the cells into micrometer-sized constrictions in a microfluidic device (**Fig. 4 A-B** and **Video S1-S2**). This microfluidic approach allowed us to measure the cell entry dynamics and derive their overall cellular mechanical parameters, which are markers of the cell ability to deform under constraint (Jebane et al., 2023). In a first analysis, we measured the cell entry time in the constrictions (**Fig. S4A**): SETDB1 KO cells, which show H3K9me3 enrichment at LADs, displayed significantly prolonged entry times compared to control A549 cells, suggesting an increased cell viscosity.

**Figure 4.**
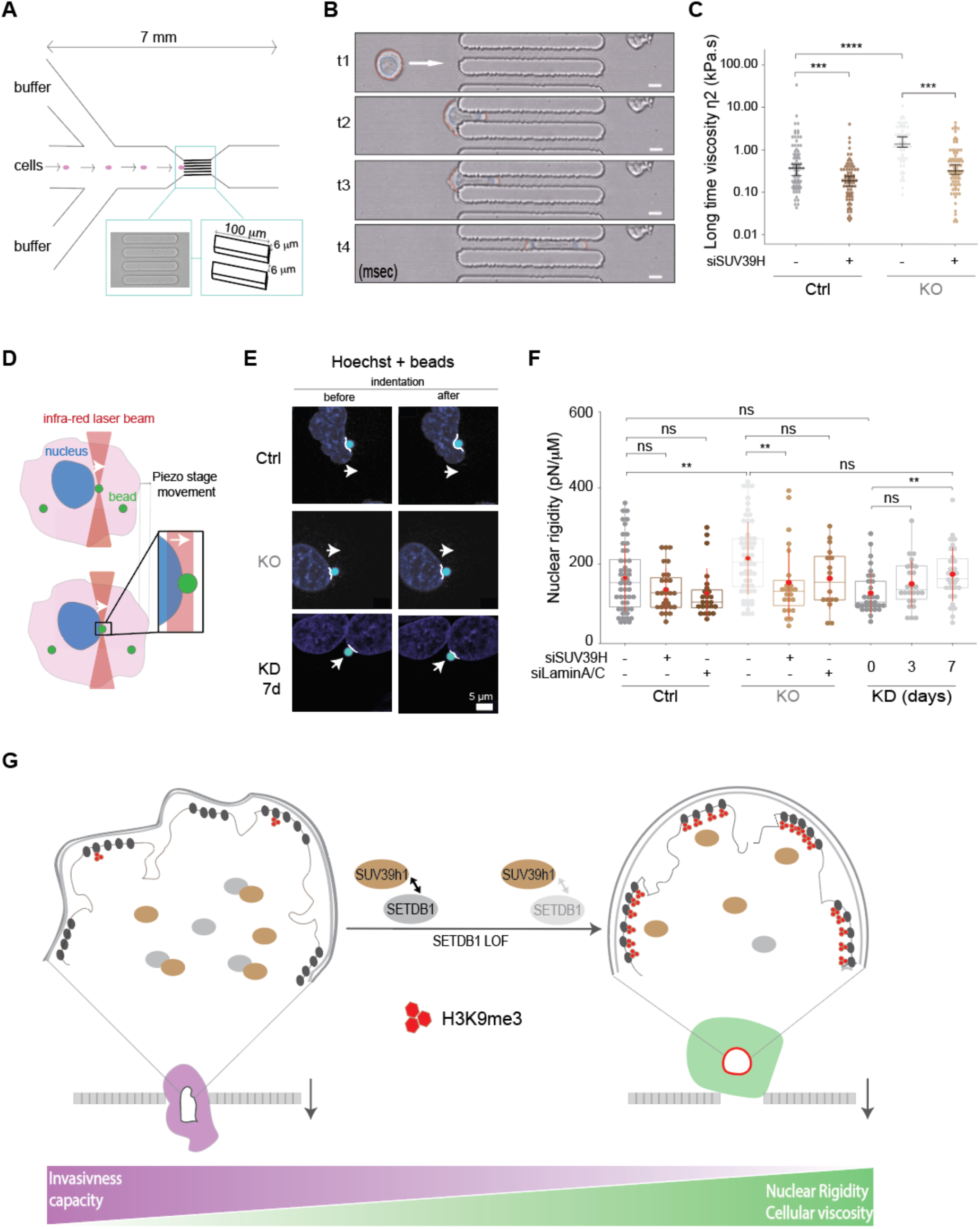
SUV39H1-induced H3K9me3 increase at LADs impacts cellular viscosity and the nuclear rigidity. **A.** Schematic representation of the microfluidic assay. Cells are pushed under controlled pressure drop through constrictions of square cross-section (6×6 μm^2^) and the entry of the cell was observed using brightfield microscopy at high frame rate. **B.** Typical sequential images of a cell passing through a microfluidic constriction at 4 time points (msec). **C.** Dot plot representation of the cellular long-time viscosity h_2_, in kPa.s, of Ctrl cells and SETDB1 KO cells with siCTR and siSUV39H. The dot plots of the non-Gaussian distributions are displayed in log scale; each dot corresponds to one cell; medians and 95% CI are indicated. Data represent N = 3 biological replicates, each with more than 50 cells (n > 50). Statistical significance was determined using the Wilcoxon test; ns indicates p > 0,05, * indicates p < 0.05, ** indicates p < 0.01, *** indicates p < 0.001, **** indicates p < 0.0001. **D.** Schematic representation of the optical tweezers nuclear indentation experiment. After internalization of 2 µm-diameter green fluorescent beads by the cells, a bead located close to the nucleus is trapped by the optical tweezers. The piezo stage is moved in order to push the bead against the nuclear envelope and indent the nucleus. **E.** Sequential images of a nucleus before and after the piezo stage movement and the indentation for Ctrl, SETDB1 KO, SETDB1 KD 7d cells. The nuclei were stained in blue with Hoechst; the bead is shown in green. **F.** Boxplot showing the quantification of the nuclear rigidity K (pN/μm) in Ctrl and SETDB1 KO cells treated with siCTR, siSUV39H, siLaminA/C and in SETDB1 KD 0, 3d and 7d dox treated cells. The line that divides the box represents the median of the data. The ends of the box indicate the upper (Q3) and lower (Q1) quartiles. The difference between Quartiles 1 and 3 is known as the interquartile range (IQR), which measures the spread of the middle 50% of the data. The lines extending from the box show the range of values within Q3 + 1.5 × IQR to Q1 - 1.5 × IQR, representing the highest and lowest values, excluding outliers. Dots beyond the whiskers indicate potential outliers in the dataset. The mean +/− SD is shown in red. Statistical significance was determined using the Wilcoxon test; ns indicates p > 0,05, * indicates p < 0.05, ** indicates p < 0.01, *** indicates p < 0.001, **** indicates p < 0.0001. **G.** Working model for the regulation of H3K9me3 distribution and nuclear mechanics by SETDB1 and SUV39H1. The balance between SETDB1 and SUV39H1 determines the localization of H3K9me3 at LADs. Upon SETDB1 loss of function (LOF), SUV39H1 activity drives the redistribution and enrichment of H3K9me3 at the nuclear periphery. Increased H3K9me3 at LADs enhances cellular and nuclear biophysical properties: it induces an increase of cellular viscosity and nuclear stiffness, reducing nuclear deformability and impairing efficient cell invasion.

Because cell size also influences the passage through the microfluidic constrictions, we assessed whether cell size contributed to these differences. Indeed, SETDB1 KO cells were larger than control A549 cells and both SUV39H KD cells (**Fig. S4B**). Importantly, sorting cells by size did not alter the observed trends in entry time across the different conditions (**Fig. S4C**), suggesting that the slowing down of the SETDB1 KO cells is due to an altered structure of the cell and its components, rather than altered size.

We next performed a finer analysis of the entry dynamics and extracted three cellular mechanical parameters using the Jeffreys viscoelastic model (see Materials and Methods): the elastic modulus E, the short-time viscosity □_1_ (which reflects the viscosity at short time scale and small deformations) and the long-time viscosity □_2_ (which reflects the viscosity at longer time scale and larger deformations). Both viscosities show a significant increase (more than twofold) in SETDB1 KO cells compared to A549 control cells (**Fig. 4C** and **Fig. S4D**). These viscosities describe the rate of cellular deformation and their increase is consistent with the observed prolonged entry time. On the other hand, the elastic modulus E slightly decreased, though not significantly (**Fig. S4E**), indicating little change of the cell overall stiffness.

Knocking down SUV39H via siRNA affected the cell mechanics of A549 control and SETDB1 KO cells in a similar way, with an important decrease of both viscosities, consistent with a decreased entry time, and a slight increase of the elastic modulus (**Fig. 4C** and **Fig. S4 A-E**). Our microfluidic experiments show that STEBD1 KO and SUV39H KD have opposite effects on the cellular overall viscosity, similarly to their opposite effects on H3K9me3 localization at LADs. These results suggest that cellular viscosity is sensitive to H3K9me3 distribution, in particular at LADs.

Since microfluidic measurements reflect only the overall mechanical properties of the whole cell, we next sought to specifically assess nuclear envelope rigidity. To determine whether H3K9me3 levels at LADs influence nuclear envelope rigidity in cancer cells, we used optical tweezers to deform the nuclear envelope and quantify nuclear stiffness (**Fig. 4D**). We found that nuclei from control A549 cells were more deformable than those from A549 SETDB1 KO cells, which exhibit increased H3K9me3 enrichment at LADs (**Fig. 4 E-F** and **Video S3-S4**). Conversely, reducing H3K9me3 levels at LADs via SUV39H KD decreased nuclear stiffness (**Fig. 4F**), a known model of altered nuclear architecture (Hah & Kim, 2019). Similarly, shRNA-mediated partial KD of SETDB1, which induces H3K9me3 enrichment at LADs without major transcriptional changes, led to increased nuclear rigidity. After 7 days of shRNA induction, nuclear stiffness in SETDB1 KD cells approached the levels observed in SETDB1 KO cells (**Fig. 4 E-F**), strongly suggesting that the H3K9me3 increase at the LADs alone is sufficient to impact the physical properties of the nucleus independently of transcriptional changes.

We also noticed that increased nuclear rigidity was associated with more circular nuclear morphology and that the proportion of circular nuclei was higher in both KO and 7d KD cells compared to controls (**Fig. S4F**). Although SETDB1 KO cells showed a stronger trend toward correlation between nuclear circularity and rigidity, this relationship did not reach statistical significance (**Fig. S4G**). These findings suggest that the observed increase in nuclear rigidity is primarily driven by H3K9me3 enrichment at LADs, rather than changes in nuclear shape alone. Conversely, increased circularity may be a consequence of H3K9me at LADs and of the subsequent increase in nuclear rigidity.

Collectively, these results demonstrate that elevated H3K9me3 levels at LADs increase both cellular viscosity and nuclear rigidity. Although these changes are accompanied by modifications in cell size and nuclear circularity, those morphological parameters alone cannot account for the observed biophysical changes, as their correlations did not reach statistical significance. Notably, in the KD model, where SETDB1 is only partially reduced and gene expression remains largely unchanged (**Fig. 1 C-F**), the observed impaired invasiveness capacities (**Fig. 1H**) could be attributed to increased H3K9me3 at LADs, which stiffens the nucleus and increases cell viscosity.

In summary, the balance between SETDB1 and SUV39H1 determines the distribution of H3K9me3 at LADs: when SETDB1 is depleted, H3K9me3 is redistributed and enriched at the nuclear periphery (LADs), a process largely driven by SUV39H1 activity. This redistribution of H3K9me3 does not broadly change LAD genomic organization but alters cellular and nuclear biophysical properties. Specifically, increased H3K9me3 at LADs enhances cellular viscosity and nuclear rigidity, making cells less able to deform and impairing their efficient migration (**Fig. 4G**). Our findings suggest that heterochromatin structural changes, particularly H3K9me3 enrichment at LADs, play a key role in controlling nuclear mechanics and limiting cell motility.

## DISCUSSION

Our study reveals an unexpected role for the histone methyltransferase SETDB1 in modulating the nuclear distribution of H3K9me3 and, consequently, the mechanical properties and migratory behavior of cancer cells. Although SETDB1 is primarily known for depositing H3K9me3 and repressing gene transcription, we show that its overexpression in lung cancer cells paradoxically prevents the accumulation of H3K9me3 at LADs. We suggest a potential mechanism in which the overexpressed SETDB1 may perturb SUV39H1 localization and partially sequester it, limiting the availability of this key H3K9 tri-methyltransferase at the nuclear periphery (**Fig. 4G**). Upon SETDB1 depletion, SUV39H1 redistributes and deposits H3K9me3 at the nuclear lamina. This epigenetic reorganization increases nuclear stiffness and cell viscosity, ultimately impairing migration.

Previous studies demonstrated the dual role of heterochromatin: it impacts both transcriptional reprogramming and biophysical properties during migration (Gerlitz, 2020). To uncouple effects on biophysical properties from effects on transcriptional changes, we induced a partial SETDB1 reduction that impairs migration without significant alterations in gene expression. Our data establish a mechanistic link between epigenetic architecture and cellular biophysics, showing that chromatin organization, specifically H3K9me3 localization, can regulate nuclear mechanics and, consequently, cell motility independently of transcriptional changes.

Other studies have observed that heterochromatin organization responds dynamically to mechanical stimuli. For instance, mechanical stretch leads to nuclear deformation and a reduction in H3K9me3 levels, potentially to prevent DNA damage (Nava et al., 2020). Similarly, melanoma cells subjected to repeated passages through narrow constrictions exhibit persistent changes in 3D genome architecture along with heterochromatin redistribution, Lamin A/C rearrangement, and increased migratory capacity (Golloshi et al., 2022). While previous studies show that chromatin changes occur after mechanical stimulation, our results reveal that pre-existing chromatin states can shape nuclear mechanics and cell migration. This is especially relevant in cancer, where epigenetic regulators like SETDB1 are often dysregulated. Notably, in our model, H3K9me3 redistribution upon SETDB1 loss leads to a characteristic ring-like pattern at the nuclear periphery, similar to that seen in non-cancerous epithelial cells (Zakharova et al., 2022). This reorganization is entirely dependent on the enzymatic activity of SETDB1, as only the wild-type protein and not the catalytically inactive form, restores the cancer-specific H3K9me3 distribution. Notably, this shift occurs without major changes to the identity of LADs themselves. An exception is chromosome 19, where LAD sequences differ, likely because many of its sequences are direct SETDB1 targets.

The relevance of chromatin-mediated nuclear mechanics extends beyond cancer. For example, recent work demonstrated that in endothelial cells, pulsatile shear stress induces SUV39H1-dependent enrichment of H3K9me3 at the nuclear periphery through interactions with Emerin, a lamina-associated protein (Chen et al., 2025). This leads to repression of inflammatory genes and increased nuclear stiffness, providing protection during blood flow. These findings underscore the context-dependent role of nuclear stiffness: while increased rigidity can be protective in the vasculature, it may hinder migration during certain steps of the metastatic cascade. For instance, successful passage through tight epithelial or endothelial barriers likely requires a more deformable nucleus with reduced heterochromatin, whereas survival in the circulation may benefit from a stiffer, more protective chromatin state.

While A549 cells serve as a well-characterized model for NSCLC, they may not fully capture the heterogeneity of lung cancer *in vivo*. Thus, validating our findings in additional NSCLC cell lines and primary patient-derived tumor samples will be important. Moreover, other H3K9 KMTs (e.g., G9a/GLP or SETDB2), other epigenetic modifications (such as DNA methylation), or chromatin-associated proteins such as HP1 and components of the nuclear lamina may also contribute to the regulation of LAD-localized chromatin states and nuclear mechanics.

This work opens several avenues for future investigation. First, dissecting the precise molecular interplay between SETDB1 and SUV39H1 may reveal biomarkers or epigenetic signatures in highly metastatic cells. Second, our observation that impaired migration can occur in the absence of major transcriptional changes challenges the traditional expression-centric view of epigenetic regulation. This points to a broader paradigm in which the mechanical and structural roles of chromatin must be integrated into our understanding of cellular behavior and fate. A valuable extension of this study would be a complementary analysis of other cellular components, particularly the cytoskeleton, to gain a more comprehensive understanding of how LAD reorganization impacts overall cell mechanics. Previous work has shown that interactions between the nucleus and the cytoskeleton play a role in changing cellular mechanical properties (Guo et al., 2014; Jebane et al., 2023). Finally, our findings can have implications beyond cancer biology, as nuclear rigidity, LAD-associated chromatin modifications, and SUV39H1 activity could also be relevant to aging, cellular differentiation, and laminopathies.

In summary, this study uncovers an unexpected role for SETDB1 in limiting H3K9me3 accumulation at LADs in cancer cells through its functional interplay with SUV39H1. Our results establish a mechanistic link between chromatin architecture, nuclear mechanics, and cell migration. Together, these findings provide a foundation for future investigations into how epigenetic regulators drive metastatic progression through both transcriptional and non-transcriptional mechanisms.

## Supporting information

Microfluidics.A549Ctrl

Microfluidics.SETDB1KO

OpticalTweezers.A549

OpticalTweezers.SETDB1KO

## RESOURCE AVAILABILITY

## Lead contact

Further information and request for resources and reagents should be directed to the lead contact, Dr Slimane Ait-Si-Ali.

## Materials availability

Materials generated in this study are available from the lead contact upon request.

## Data availability

- The DAM-ID and RNAseq data that support the findings of this study are available under the accession numbers XXXX and XXXX.
- All data needed to evaluate the conclusions in the paper are present in the paper and/or the Supplementary Materials.
- Further details necessary for reanalysis of the data presented in this paper are available from the lead contact upon request.

## AKNOWLEDGMENTS

Work in the Ait-Si-Ali lab was supported by the Agence Nationale de la Recherche (ANR, ANR-24-CE45-0690-03 grant to S Ait-Si-Ali J.B. Manneville and E. Helfer), Association Française contre les Myopathies Telethon (AFM-Telethon, grant # 22480, to S Ait-Si-Ali); Fondation pour la Recherche Médicale (FRM, « Equipe FRM » grant # DEQ20160334922, to S Ait-Si-Ali); Université Paris Diderot (now Université Paris Cité) through the “Who Am I?” Laboratory of Excellence and the Emergence call (2024-ChromaStiff grant), # ANR-11-LABX-0071, to S Ait-Si-Ali, funded by the French Government through its “Investments for the Future” program, operated by the ANR under grant #ANR-11-IDEX-0005-01. S.C. was supported by PhD fellowships from the doctoral school BioSPC of Paris Cité University and Fondation ARC pour la Recherche sur le Cancer.

CJ was supported by Excellence Initiative of Aix-Marseille University - A*MIDEX (A-M-AAP-ID-17-66-170301-11.30) funding to E. Helfer and by ANR grant ANR-17-CE12-0010 to S. Ait-Si-Ali. PL is supported by the Agence Nationale de la Recherche (ANR, ANR-24-CE45-0690-03 grant to E. Helfer, S Ait-Si-Ali, and J.-B. Manneville). EH belongs to the French Consortium Approches Quantitatives du Vivant/Quantitative approaches to living systems (GDR AQV). G. Lamour is supported by ANR grant N° ANR-21-CE19-0027–04 and the University of Evry-Paris Saclay. J.-B. Manneville is supported by by grants from ITMO Cancer Inserm-Aviesan “Approches interdisciplinaires des processus oncogéniques et perspectives thérapeutiques : Apports à l’oncologie de la physique, de la chimie et des sciences de l’ingénieur” Edition 2022” (NUTMEG project, grant #22CP073-00) and from the Labex “Who Am I?” (ANR-11-LABX-0071) and the “Initiatives d’Excellence” (Idex ANR-11-IDEX-0005-02) transverse project BioMechanOE (TP5).

We thank Tara Stoll-Bickel for the technical help in the western blot assays. We thank the (EPI)2 Imaging platform (managed by Sandra Piquet and Audrey Chansard) - UMR7216 Epigenetic and Cell Fate Centre, for access to instruments and technical advice, with financial support from ITMO Cancer of Aviesan within the framework of the 2021-2030 Cancer Control Strategy, on funds administered by Inserm. We thank the GENIE platform, Epigenetic and Cell Fate center, for technical advice and practical help. We thank members of the Ait-Si-Ali lab and Epigenetics and Cell Fate department for helpful discussions during the group and department meetings and critical reading of the manuscript.

## AUTHOR CONTRIBUTIONS

Conceptualization, S.A., E.B.; Methodology Software; G.V., M.H.; Formal Analysis; S.C., V.P., E.B., L.D., G.V., V.Z., C.J., P.L., V.J.; Investigation; S.C., V.P., E.B., V.Z., C.J., P.L.; Data Curation, S.C., V.P., E.B., G.V., C.J., P.L., E.H., J.B.M; Writing – Original draft; S.C.; Writing – Review & Editing; S.A., E.B., S.C., G.L., E.H., J.B.M., G.V., V.J., P.L., C.J., V.Z.; Supervision, S.A.; Funding Acquisition, S.A., E.H., J.B.M; Project Administration, S.A.

## MATERIAL AND METHODS

### Cell culture conditions

A549 human male lung carcinoma cells (CCL-185) were obtained from ATCC and cultured in Dulbecco’s Modified Eagle’s Medium (DMEM; Gibco, ref. 31966-021 DMEM 1X + Glutamax^TM^-I), supplemented with 10% fetal bovine serum (FBS; Dutscher, ref. 500105M1M) and 1% penicillin/streptomycin (Gibco, ref 15140-122). Cells were maintained at 37°C in the presence of 5% CO_2_ and were periodically screened for *Mycoplasma* contamination.

### Cell lines establishment

Stable shSETDB1-A549 cell lines were established by transducing cells with in-house–produced lentiviral particles carrying the constructs listed in Table 1, cloned into pLKO-Tet-On vectors (modified from Addgene#83481). shRNA sequences targeting SETDB1 and a scrambled shRNA used as a negative control were designed using the siRNA Wizard tool (https://www.invivogen.com/sirnawizard/). shRNA expression was induced with doxycycline (1 µg/mL). The vector contains two antibiotic resistance genes: ampicillin (for bacterial selection) and puromycin (for selection of stable integration in eukaryotic cells).

**Table 1.**
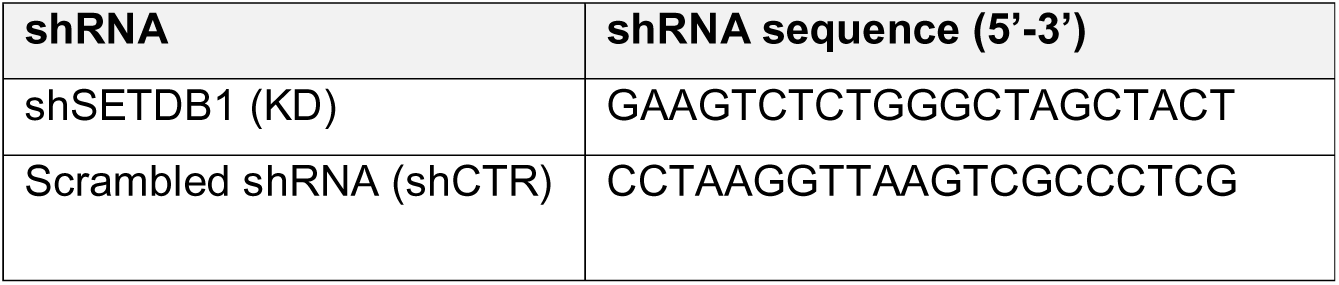
SETDB1 and scrambled shRNA sequences.

A549 SETDB1 KO clone (#32) was obtained by inducing the complete loss of SETDB1 expression using CRISPR/Cas9, as described in (Zakharova et al., 2022). To generate SETDB1 KO cell lines expressing either wild-type (SETDB1 WT) or mutant forms (hSETDB1H1224K; K1170A+K1178A) for the rescue experiments (**Fig. 3 C-D**), SETDB1 KO cells were transduced with in-house–produced lentiviral particles containing a modified version of the pLKO-Tet-On vector (derived from Addgene #83481). Polyclonal populations were subsequently selected using Blasticidin S HCl (10 μg/ml; Gibco, ref. A11139-03). To generate A549 cell lines expressing either wild-type or mutant (H324K) forms of SUV39H1 (**Fig. 3 A-B**), A549 cells were transduced using the procedure, followed by selection with Hygromycin B (50mg/mL, Invitrogen, ref. 20687010). The plasmids used in this study are listed in **Table 2**. The indicated modifications were introduced using the NEBuilder HiFi DNA Assembly Master Mix (NEB, E2621S). Cloning primers were designed using the NEBuilder Assembly Tool (https://nebuilder.neb.com/), and site-directed mutagenesis was performed using the Q5 Site-Directed Mutagenesis Kit (NEB, E0552S)

**Table 2.** List of plasmids.

| Insertion | Plasmid sequence |
| --- | --- |
| Control | pCW_TetON_mCherry |
| SETDB1 WT | pCW_TetOn_Flag_HA_hSETDB1-P2A-mCherry |
| SETDB1 hSETDB1H1224K | pCW_TetOn_Flag_HA_hSETDB1H1224K-P2A-mCherry |
| SETDB1 mut K1170A+K1178A | pCW blast-flag_HA_hSETDB1mut_bgh polyA_hPGK_BLST_T2A _reverse tetracycline-controlled transactivator |
| SUV39h1wt | pCW_Myc_hSuv39h1_bgh polyA_hPGK_Hygro_T2A _reverse tetracycline-controlled transactivator |
| SUV39h1 mut H324K | pCW_Myc_hSuv39h1_H324K_bgh polyA_hPGK_Hygro_T2A _reverse tetracycline-controlled transactivator |

**Table 3.**
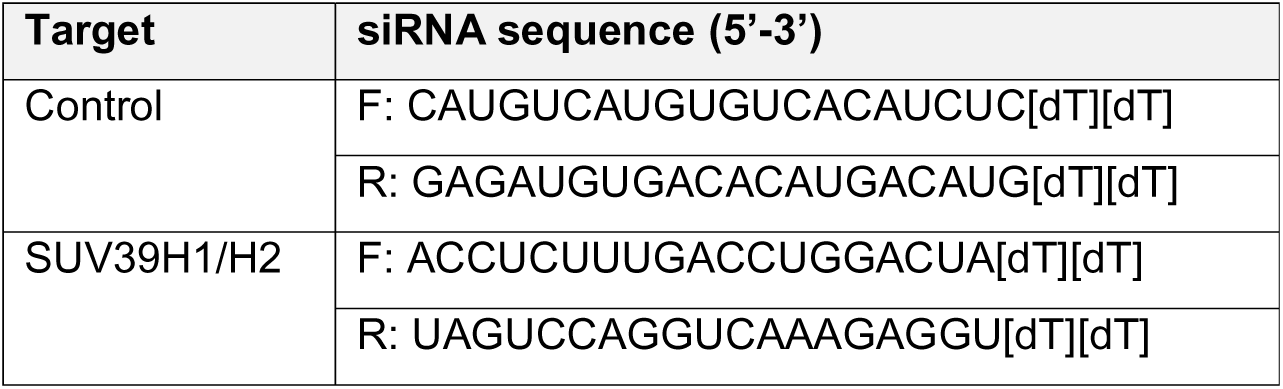
List of siRNA sequences.

### Gene silencing by siRNA

siRNAs targeting human SUV39H (SUV39H1/H2) and a non-targeting negative control siRNAs were obtained from Sigma-Aldrich. siRNAs were diluted with Opti-MEM® Reduced Serum Medium (Gibco, 1X, ref. 11058-021). Transfections were performed at final concentration of 20 nM siRNAs using the Lipofectamine® RNAiMAX Reagent (Invitrogen™, ref. 13778150). Cells were harvested 72h post-transfection. The efficiency of gene silencing was evaluated by western blotting and RT-qPCR.

### Immunofluorescence

For immunofluorescence imaging, cells were seeded on glass coverslips at 60–70% confluence, fixed in 2% paraformaldehyde (PFA) and permeabilized with PBS containing 0.2% Triton X-100. Samples were blocked with 3% BSA in PBS-T, then incubated overnight at 4◦C with appropriate primary antibodies (**Table 4**). After three washes three with PBS-T, samples were incubated for 1h at room temperature with appropriate fluorophore-conjugated secondary antibodies (anti-rabbit 1:400 Alexa 594 IgG; anti-mouse 1:400 Alexa 488; Jackson ImmunoResearch Laboratories). Cell nuclei were stained with DAPI (Life Technologies; Cat#: 62248) and coverslips were mounted using Vectashield mounting media (Clinisciences).

**Table 4.** Antibodies list.

| Antibody | Reference | Use (dilution) |
| --- | --- | --- |
| H3K9me3 | Diagenode; Cat#: C1541093 | IF (1:500) |
| E-cadherin | BD Biosciences, 610181 | IF (1:1000) |
| phalloidin-TRITC | Sigma-Aldrich; Cat#: P1951 | IF (1:1000) |
| SETDB1 | Proteintech, 11231-1-AP | Western blot (1:1000) |
| Lamin A/C | Proteintech, 10298- 1-AP | Western blot (1:1000); IF (1:100) |
| FLAG tag | Sigma-Aldrich, F3165 | IF:1/1000 |
| MYC tag | Abcam, ab32 | IF:1/1000 |
| SUV39h1 | CST, 8729 | Western Blot (1:200) |
| H3 | Santa Cruz Biotechnology, sc-8654 | Western Blot (1:1000) |
| Ezh2 | CST, 3147 | Western Blot (1:1000) |

Imaging was performed using an inverted Leica DMI-6000 microscope equipped with a 100× oil-immersion objective. Images were acquired using the HQ2 Coolsnap camera controlled by MetaMorph 7.10.2.240 software. All images were processed with ImageJ (Fiji) software.

For automated imaging with the PerkinElmer® Operetta CLS high-content microscope, cells (2.6×10^4^) were seeded directly onto a 96-well plate and fixed with 2% PFA after 24h. Staining was performed as described above. Images were acquired using Andor Zyla 5,5 sCMOS 16 bit 5,5 Mpx camera and a 40X water-immersion objective (NA 1.1; WD: 0.62 nm), with a pixel size (binning 1) of 0.1625 μm and a maximum resolution of 330 nm at 488. The border area (+/− 10 % distance from the periphery) was defined based on the DAPI staining imaged with the Operetta software. The H3K9me3 intensity at the nuclear periphery was calculated as the ratio of border H3K9me3 signal/ - H3K9me3 signal. Three replicates were included in the analysis and statistical significance was determined using the Wilcoxon test; ns indicates p > 0,05, * indicates p < 0.05, ** indicates p < 0.01, *** indicates p < 0.001, **** indicates p < 0.0001.

### Nuclei fractionation

Cells were counted and lysed at a concentration of 2 × 10□ cells/μL in Buffer A (20 mM HEPES, pH 7.0; 0.15 mM EDTA; 0.15 mM EGTA; 10 mM KCl) containing 10% NP-40 and SR buffer (50 mM HEPES, pH 7.0; 0.25 mM EDTA; 10 mM KCl; 70% (m/v) sucrose), supplemented with protease inhibitors (Sigma) and spermidine/spermine (0.15 mM each) to prevent nuclear leakage. Cell lysates were centrifuged at 2,000 × g for 5 min, and the resulting nuclear pellets were resuspended in low-salt buffer (20 mM Tris, pH 7.65; 0.2 mM EDTA; 25% glycerol; 1.5 mM MgCl_₂_; 15 mM KCl) to a final concentration of 1 × 10□ nuclei/ μL. For each extraction condition, 30 μL aliquot of the nuclear suspension (3 × 10□ nuclei) was used per extraction condition.

An equal volume of high-salt buffer (20 mM Tris-HCl, pH 7.65; 0.2 mM EDTA; 25% glycerol; 1.5 mM MgCl_₂_) containing NaCl was added to achieve final NaCl concentrations ranging from 400 mM to 1 M. Samples were incubated for 1 h on ice, then centrifuged at 16,000 × g for 90 at 4 °C. The soluble fraction was collected for subsequent western blotting analysis. For total nuclear samples, after incubation on ice, the suspension was sonicated for 5 min (15 s on/ 15 s off) at high frequency using a Bioruptor (Diagenode) and used directly for western blotting preparation. Extracts were diluted up to a final NaCl concentration of 300mM, supplemented with NuPAGE LDS sample buffer (1x) and 100 mM DTT. Equal volumes of each extract were loaded for analysis.

### Western blot

Cells were counted, and the pellet resuspended in loading buffer (NuPAGE LDS sample buffer 1x, NuPAGE sample reducing agent 1w, 10mM Tris-HCL pH 8.0), then boiled for 5min at 95°C. Proteins were separated on NuPAGE 4–12% Bis-Tris polyacrylamide gels (Invitrogen) using 1× NuPAGE MOPS SDS running buffer, and transferred into nitrocellulose membrane (Sigma-Aldrich, Amersham, ref. 10600002) in 20 mM phosphate transfer buffer (pH 6.7). Membranes were blocked in 5% milk in PBS-T buffer (1× PBS, 0.2% Tween 20) and incubated overnight at 4°C with the appropriate primary antibody (Table 4). After two 5-min washes in PBS-T, membranes were incubated with the appropriate IRDye secondary antibodies (LI-COR) in PBS-T, followed by two 10-min washed twice in PBS-T, and one 10 min wash in PBS. Signals were visualized using the Odyssey Imaging System (LI-COR).

### RNA extraction and quantification, RT-qPCR

Total RNA was extracted using the RNeasy® Plus Mini Kit (QIAGEN, cat# 74106) according to the manufacturer’s instructions, with automated processing on the QIAcube system (QIAGEN). cDNA synthesis was carried out using MultiScribe^TM^ Reverse Transcriptase (ThermoFisher) following the manufacturer’s instructions. Quantitative PCR reactions were prepared with SYBR™ Green reagent (Applied Biosystems®) using a TECAN pipetting robot, and amplification was performed on a Pro6 real-time PCR system (Applied Biosystems®). Primers were synthesized by Sigma Aldrich (**Table 5**). Relative mRNA expression levels were calculated using the ΔΔCt method with *PPIA* (*Peptidylprolyl Isomerase A*) serving as the internal normalization control

**Table 5.**
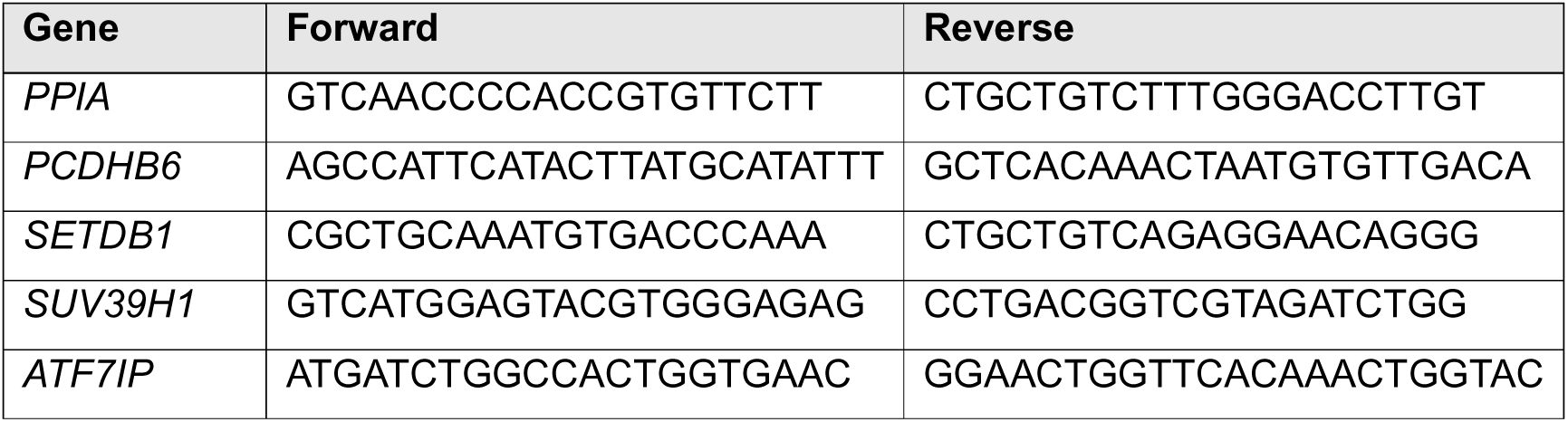
List of primers sequences.

### RNA-sequencing

Total RNA was isolated using RNeasy Mini Kit (Qiagen; Cat#: 74104), followed by treatment with the Turbo DNA-free Kit (Thermo Fisher; Cat#: AM1907) to remove genomic DNA contamination. RNA integrity was verified (RIN > 9.9), and libraries were sequenced on an Illumina Nextseq 500 platform in paired-end mode using three independent biological replicates at <u>Platform GENOM’IC -</u> <u>Institute Cochin (Paris, France)</u>.

### RNA-seq Analysis

RNA-seq data were processed using the RASflow_EDC v1.3 pipeline (<u>BiBs-EDC</u>),, an adaptation of the original RASflow workflow by <u>X. Zhang</u>. Full documentation for RASflow_EDC v1.3 is available at: https://parisepigenetics.github.io/bibs/edctools/workflows/rasflow_edc. Reproducibility among biological replicates was assessed using MultiDimensional Scaling (MDS) and Principal Component Analysis (PCA) of expression profile distances. Differential expression analysis generated tables of differentially expressed genes (DEGs) for each comparison. Genes with an absolute log2 fold-change ≥ 0.5 were subjected to Gene Ontology (GO) enrichment analysis using the PANTHER classification system (version 19.0, released 2024-06-20; https://geneontology.org), under the GO-Slim Biological Processes category. Genes associated with the GO term Locomotion (GO:0040011) were selected to heatmap visualization. Additionally, genes related to cell migration were analyzed without applying a log_₂_FC cutoff. For this analysis, genes annotated under GO Biological Process Cell migration (GO:0016477) were retrieved from <u>AmiGO 2</u>, and a heatmap was generated based on this gene set. Gene Set Enrichment Analysis (GSEA) was performed on count tables generated by RASflow_EDC v1.3. using GSEA software v4.3.3 with 1000 permutations. The Hallmark gene set collection (<u>h.all.v2024.Hs.symbols.gmt) was used for</u> enrichment analysis

### Cell proliferation

Cells (0,5×10^5^) were seeded in a 6-well plate. After 3, 5 and 7 days, cells were counted using an automated cell counter. Data are presented as mean□±□SD. Three independent replicates were included in the analysis and statistical significance was determined using an unpaired t-test; ns indicates p > 0,05, * indicates p < 0.05, ** indicates p < 0.01, *** indicates p < 0.001.

### Wound-healing assay

Cells were seeded in 12-well plates and cultured for 24□h to reach 90 to 100% confluence. After washing with PBS, a sterile 200□μL pipette tip was used to make a perpendicular scratch across the cells’ monolayer. Detached cells were removed by washing with PBS and fresh medium added. Cells were then incubated for 48h. The scratch closure was monitored and imaged at 0h (immediately after scratching), at 24h and at 48h using a live-cell microscope with 5x objective. Scratch closure was analyzed using Fiji (ImageJ) software by measuring the wound width at five different positions per image, calculating the average percentage of wound reduction over time. Data are presented as mean□±□SD. Three replicates were included in the analysis and statistical significance was determined using an unpaired t-test; ns indicates p > 0,05, * indicates p < 0.05, ** indicates p < 0.01, *** indicates p < 0.001.

### Transwell assay

Cells (3×10^4^) were seeded into Transwell chambers containing polycarbonate membrane with 8 mm pores (Corning ref. 3422). After 24h, migrated cells on the lower surface of the membrane were stained with Crystal Violet, and images were taken using a live-cell microscope. For quantification, Crystal Violet was solubilized with 33% acetic acid, incubated for 10 min on a shaker, and the absorbance was measured at 590 nm using a plate reader to quantify migrated cells. Data are presented as mean□±□SD. Three replicates were included in the analysis and statistical significance was determined using an unpaired t-test; ns indicates p > 0,05, * indicates p < 0.05, ** indicates p < 0.01, *** indicates p < 0.001.

### Nuclear indentation with optical tweezers

Cells were plated on WPI Fluorodish^TM^ dishes (FD35-100). 2 µm-diameter amine-coated fluorescent beads (DiagPolyTM #DCGF-L022) were added to the cell culture (1:10 000 dilution) and incubated with the cells for 72h at 37°C with 5% CO_2_. Cells were stained with 90 Hoechst (33384) at a 1:10 000 dilution in cell culture medium for 10 mins before the nuclear rheology experiment. The experimental setup combining optical tweezers with fast confocal microscopy was previously described in detail in (Guet et al., 2014). Briefly, a single fixed optical trap was generated by directing an infrared laser beam into the back port of an inverted Eclipse microscope (Nikon), equipped with a resonant laser confocal A1R scanner (Nikon), a 37□°C environmental chamber, and a nanometric piezo stage (Mad City Labs). The nuclear indentation protocol and associated analysis procedures followed (Alibert et al., 2021). In summary, to indent the nucleus, a bead in contact with the nuclear envelope was first captured by the optical trap. The piezo stage was then moved at a constant velocity, pushing the nucleus toward the trapped bead. The applied force was calculated based on the bead’s displacement from the trap center (trap stiffness: 240□pN/µm), while the indentation depth was determined via image analysis. Nuclear rigidity was inferred from the resulting force-indentation curves using a viscoelastic model as described by (Alibert et al., 2021). The nucleus rigidity K (pN/μM) values are represented in boxplots where the line that divides the box represents the median of the data. The ends of the box indicate the upper (Q3) and lower (Q1) quartiles. The difference between quartiles 1 and 3 is known as the interquartile range (IQR), which measures the spread of the middle 50% of the data. The lines extending from the box show the range of values within Q3 + 1.5 × IQR to Q1 - 1.5 × IQR, representing the highest and lowest values, excluding outliers. Dots beyond the whiskers indicate potential outliers in the dataset. The mean +/− SD is shown in red. Statistical significance was determined using the Wilcoxon test; ns indicates p > 0,05, * indicates p < 0.05, ** indicates p < 0.01, *** indicates p < 0.001, **** indicates p < 0.0001.

From the optical acquisitions, circularity was calculated with Fiji: 0.7 was defined as a threshold of circularity and nuclei were scored as circular if the value of circularity was greater than 0.7.

### Microfluidics

Microfluidic assays were performed as previously described in (Jebane et al., 2023).

#### Microfluidic device

Briefly, microfluidic chips in polydimethylsiloxane (PDMS) were fabricated using an house-made mold made of SU8-photoresist, prepared via photolithography (for details see: (Jebane et al., 2023)). A chip consists of a chamber made of a bottom PDMS-coated glass coverslip assembled with a few mm-thick PDMS piece with a channel dug in it (defined as “microchannel”). The 6-mm long microchannel holds 6×6 µm^2^ constrictions in the center that are used to geometrically constrain the cells passing through under a fixed pressure drop.

#### Microfluidic experiments

The experiments were performed on a microscope (IX83, Olympus) equipped with a 20× objective and high-speed cameras (Fastcam Mini UX, Photron) at 37°C. The chip inlets and outlet were connected to a pressure controller (MFCS™-EZ 1000 mbar, FLUIGENT) that allowed controlling pressures at inlets and outlet. Top and bottom inlets were used to inject culture medium and buffers while middle inlet was used to inject cells (see Fig. 3D for a schematic view of the chip): the pressures applied at inlets and outlet fixed the pressure drop that drives the cell flowing in the microchannel and constrictions. Microchannels were first passivated with 10% Pluronic®F-127 (P2443, Sigma) in PBS for 1h at RT, then rinsed with cell culture medium. 72h prior to experiments, cells were seeded in 3-cm Petri dishes and transfected twice with siRNA following the procedure described in “Gene silencing by siRNA” section. Cells were initially seeded to ensure ≈90% cell confluence on the day of the experiment. Once the chip was passivated and rinsed, cells were trypsinized, resuspended at 5.10^4^ cells/mL in culture medium + 10mM HEPES, then injected in microchannels under a pressure drop of ΔP = 165 mbar (Pinlets = 170 mbar, Poulet = 5 mbar). Cells were observed in brightfield and movies were acquired at frame rates varying from 250 to 1000 fps. At least three replicates were done for each condition.

#### Cell size analysis

Cell size was estimated by measuring the projected cell diameter D in the microchannel as a proxy before they reach the constrictions. Indeed, the device being 6µm high, the cells are squeezed in height and their projected shape is circular (observed from the bottom, see a typical image of a circular-shaped cell in **Fig. 4B**, top row): their volume is approximately that of a cylinder of diameter D and height h =6 µm, i.e., V = h × π × (D/2)^2^.

#### Cell entry time analysis

Movies were pre-processed using FIJI software to crop the region of interest and select the sequence of images for each cell. The time of cell contact with the constriction and cell finalizing its entry in the constriction were determined using a MATLAB home-made program. The cell entry time was defined as the difference between these two time points.

#### Extraction of the mechanical parameters of the cell

Cells were segmented using a modified version of the long variant of the Segment Anything Model 2 (SAM2Long github: https://github.com/Mark12Ding/SAM2Long). The progression of the cell ‘tongue’ into the constriction was tracked from the segmented images and its length was measured over time using a custom Python program. The tongue length is defined as the distance between the entrance of the microchannel and the tip of the cell tongue. The mechanical parameters were extracted as described in (Jebane et al., 2023). When entering the microfluidic constriction, cells undergo three distinct phases. The first phase is rapid and reflects a viscoelastic regime, corresponding to the initial deformation of the cell. The second phase occurs as the cell slows down, likely due to the passage of the nucleus through the constriction, and is dominated by viscous behavior. Finally, once the whole cell has entered the channel, its velocity increases again. From the first phase, we can extract the elastic modulus and short-time viscosity, while the second phase provides the long-time viscosity. To extract these three parameters, the tongue length as a function of time L(t) was fitted using the Jeffreys model (viscoelastic model). In this framework, the tongue length can be expressed as a function of the viscoelastic properties of the cell:

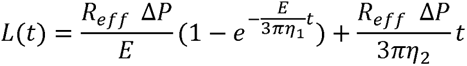

where *R_eff_* is the radius of an effective cylindrical constriction which circular cross section is equivalent in area to the square cross-section of the microchannel constrictions (6×6 µm^2^). Δ*p* is the applied pressure drop. E, *η*_1_ and *η*_2_ are the elastic modulus, the short-and long-time viscosities of the cell, respectively.

### LaminB1 DamID-seq

Cells were seeded in 6-well plates at a density of 5×10^5^ cells per well and transduced with lentiviral vectors expressing Dam or Dam-LMNB1 (Addgene plasmid #182671) in the presence of polybrene (8 µg/mL). Genomic DNA (gDNA) was extracted three days post-transduction using the Monarch Genomic DNA Purification Kit (New England Biolabs).

The DamID-seq protocol was adapted from Vogel et al. (2007) and Leemans et al. (2019). A total of 500 ng of purified gDNA was digested with DpnI (0.5 µL of 20 U/µL; New England Biolabs) in a 10 µL reaction containing 1 µL 10× CutSmart buffer, incubated at 37□°C for 5 hours, followed by heat inactivation at 80□°C for 20 minutes. DamID adapters were prepared by annealing AdRt (5′-CTAATACGACTCACTATAGGGCAGCGTGGTCGCGGCCGAGGA-3′) and AdRb (5′-TCCTCGGCCG-3′) oligonucleotides (Eurofins). Adapters were ligated to the digested DNA by adding 10 µL of ligation mix (2.5 µL T4 DNA Ligase, 1 U/µL; Roche #10716359001, 2 µL 10× ligation buffer, 0.25 µL 50 µM annealed adapter, and 5.25 µL H_₂_O) and incubating overnight at 16□°C. Ligation was followed by heat inactivation at 65□°C for 10 minutes. To remove unmethylated GATC fragments, samples were digested with DpnII (1 µL of 10 U/µL; New England Biolabs R0543L) in a reaction containing 5 µL DpnII buffer and 24 µL H_₂_O, incubated at 37□°C for 1 hour. Control reactions omitting DpnI or T4 ligase were included to evaluate amplification specificity.

Eight microliters of the digested-ligated DNA was amplified in a 40 µL PCR containing 20 µL 2× Q5 High-Fidelity Master Mix (New England Biolabs), 1 µL 50 µM Adr_PCR primer (5′-GGTCGCGGCCGAGGATC-3′), and 11 µL H_₂_O. PCR conditions were as follows: initial denaturation at 68□°C for 10 minutes; 1 cycle of 94□°C for 1 minute, 65□°C for 5 minutes, and 68□°C for 15 minutes; 5 cycles of 94□°C for 1 minute, 65□°C for 1 minute, and 68□°C for 10 minutes; followed by 18 cycles of 94□°C for 1 minute, 65□°C for 1 minute, and 68□°C for 2 minutes.

Sequencing libraries were prepared using the Illumina DNA Prep Kit starting with 100 ng of PCR product and amplified with 5 PCR cycles post-tagmentation, following the manufacturer’s instructions. Final libraries were eluted in 20 µL after bead cleanup. Library concentrations were quantified using the Qubit dsDNA HS Assay Kit, and fragment size distribution was assessed using the Agilent Bioanalyzer 2100 and TapeStation systems. Sequencing was performed on an Illumina NextSeq 2000 platform to generate an average of 50 million 1 × 100 bp single-end reads per sample. Demultiplexing and quality control were performed using Aozan software (v3.1.1) (Perrin et al., 2017).

### DamID-seq data analysis

Bioinformatic analyses were conducted using the Galaxy platform (https://usegalaxy.eu/) and RStudio (version 2024.12.1, Build 563; R version 4.4.3). Sequence quality control and trimming of DamID adapters were performed with fastp (v0.23.4). Reads were aligned to the human reference genome (hg38) using Bowtie2 (v2.5.3) with default parameters. BigWig files were generated with deepTools2 (v3.5.4) using the bamCompare function to calculate the log_₂_ ratio of normalized read counts (Dam-LMNB1 over Dam) with a bin size of 50 bp. To visualize the DamID-seq signal, bigWig files from three replicates per condition (Dam-LMNB1 and Dam) were merged and averaged using bigwigAverage (deepTools2, v3.5.4) in 100 bp bins, excluding blacklisted genomic regions. Lamina-associated domains (LADs) were identified using the HMMtBroadPeak package (Hidden Markov Model, Baum–Welch algorithm; https://github.com/jianhong/HMMtBroadPeak/). Differential LADs between A549 and A549 SETDB1 knockout (KO) cells were detected using limma (v3.58.1) with the voom transformation method. Visualization of DamID-seq data, including genomic tracks and graphical representations, was performed using the R packages *ggplot2* and *chromoMap*, alongside the Integrative Genomics Viewer (IGV).

### Statistical analyses

Statistical analyses were performed in RStudio. Data are represented as mean ± SD or median ± CI, as indicated. Two-tailed *t* test or Wilcoxon rank-sum tests were used for statistical analysis; ns indicates p > 0,05, * indicates p < 0.05, ** indicates p < 0.01, *** indicates p < 0.001, **** indicates p < 0.0001. Statistical tests used are specified in the figure legend.

## SUPPLEMENTARY MATERIAL

**Figure S1, related to Figure 1.**
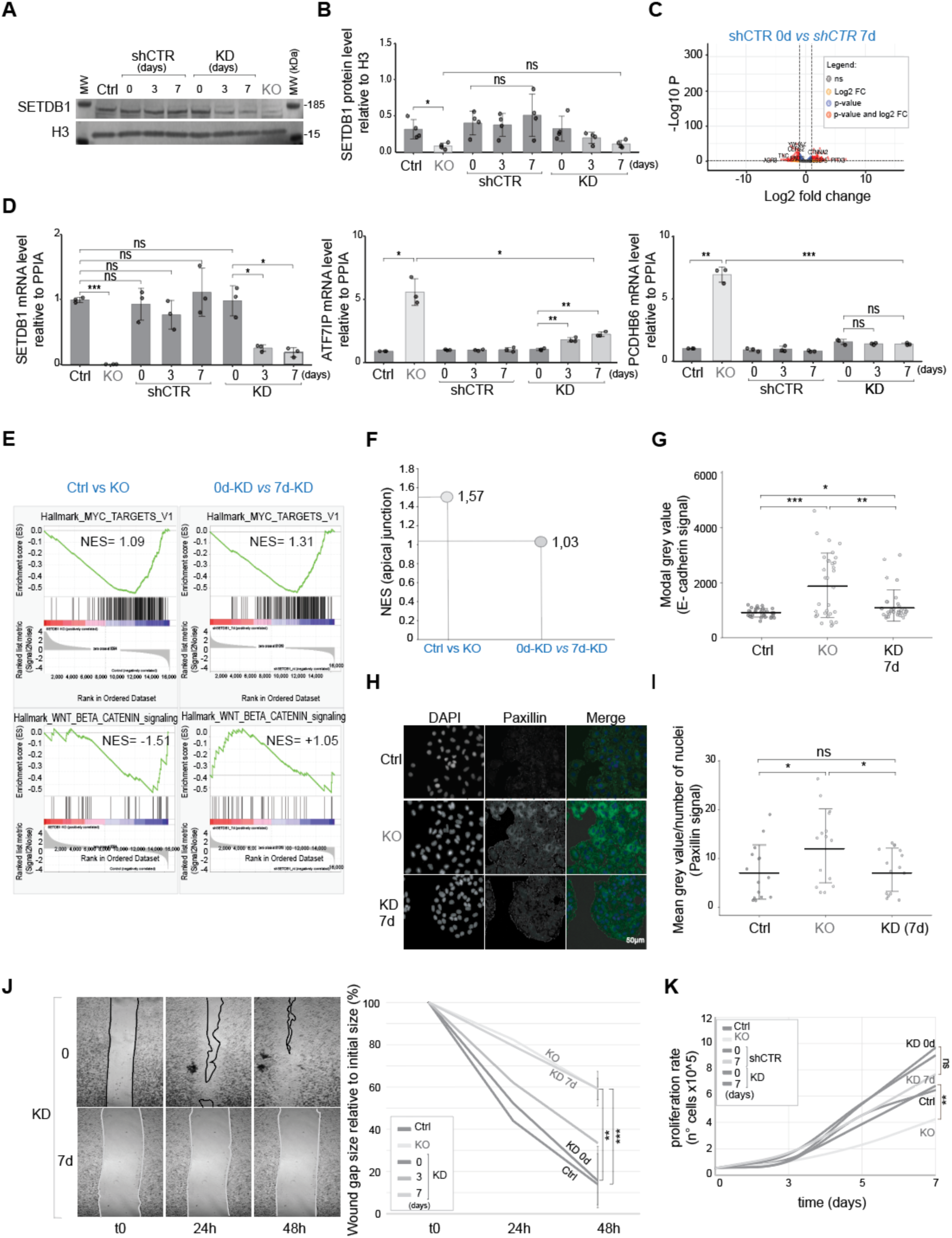
SETDB1 knockdown (KD) impairs cell migration without affecting proliferation or global gene expression, in contrast to SETDB1 knockout (KO), which alters proliferation, gene expression, and focal adhesion. **A.** Western blot analysis of SETDB1 protein levels in A549 cells (Ctrl), SETDB1 KO, shCTR (0, 3d, 7d), and SETDB1 KD (0, 3d, 7d dox-treated) cells. H3 is used as a loading control. **B.** Quantification of SETDB1 protein level normalized to H3 used for A549 cells (Ctrl), SETDB1 KO, shCTR (0, 3d, 7d), and SETDB1 KD (0, 3d, 7d dox-treated) cells. Bars represent the mean of N=4 biological replicates +/− SD error bars, with individual values shown. Statistical significance was determined using an unpaired t-test; ns indicates p > 0,05, * indicates p < 0.05, ** indicates p < 0.01, *** indicates p < 0.001. **C.** Volcano plot showing -log10(p-value) (-Log10 P) versus log2 fold change for gene expression comparison between shCTR-0d and shCTR-7d dox-treated cells, indicating minimal transcriptional effects of doxycycline or the shRNA system alone. **D.** mRNA levels of SETDB1, ATF7IP, and PCDHB6 measured by RT-qPCR and normalized to PPIA. Expression levels are relative to A549 cells (Ctrl), Bars represent the mean of N=3 biological replicates +/− SD error bars, with individual values shown. Statistical significance was determined using an unpaired t-test; ns indicates p > 0,05, * indicates p < 0.05, ** indicates p < 0.01, *** indicates p < 0.001. **E.** Gene Set Enrichment Analysis (GSEA) plots for MYC targets and WNT/β-catenin signaling pathways comparing Control vs SETDB1 KO and SETDB1 KD 0 vs 7d dox-treated conditions. Normalized Enrichment Scores (NES) are reported. **F.** NES values for the Hallmark “Apical Junction” gene set in Ctrl vs SETDB1 KO (light grey) and SETDB1 KD 0 vs 7d dox-treated (dark grey) comparisons. **G.** Quantification of E-cadherin signal (marker of cell–cell adhesion) from immunofluorescence images (see Fig. 1H) of Control, SETDB1 KO and SETDB1 KD 7d dox-treated cells. Modal grey value per field is measured, >30 fields per condition are analyzed, N= 3 biological replicates, in the graph is indicated the mean +/− SD error bars; Statistical significance was determined using an unpaired t-test; ns indicates p > 0,05, * indicates p < 0.05, ** indicates p < 0.01, *** indicates p < 0.001. **H.** Representative immunofluorescence images of Ctrl, SETDB1 KO and SETDB1 KD 7d dox-treated cells stained with anti-Paxillin (green). Nuclei are counterstained with DAPI (blue). Merged images show colocalization. Scale bar: 50 µm. **I.** Quantification of Paxillin signal (marker of focal adhesions) from images in panel S1H. Dots stand for the mean grey value of the field / number of nuclei x field, 15 fields are analyzed from N= 3 biological replicates, in the graph is indicated the mean +/− SD error bars; Statistical significance was determined using an unpaired t-test; ns indicates p > 0,05, * indicates p < 0.05, ** indicates p < 0.01, *** indicates p < 0.001. **J.** Left: Representative wound healing images at 0, 24, and 48 hours after wound for SETDB1 KD 0 and 7d dox-treated cells. Lines on pictures indicate wound edges over time Right: Quantification of wound gap size (%) over time for Ctrl, SETDB1 KO, and SETDB1 KD (0, 3d, 7d dox-treated). Wound size is normalized at time 0 (100%). Error bars stand for −/+ SD and statistical significance was determined using an unpaired t-test; ns indicates p > 0,05, * indicates p < 0.05, ** indicates p < 0.01, *** indicates p < 0.001. **K.** Cell proliferation curves showing the number of cells (×10□) over time (0, 3, 5, 7 days) for Ctrl, SETDB1 KO, shCTR (0 and 7d dox-treated), and SETDB1 KD (0 and 7d dox-treated). Statistical significance was determined using an unpaired t-test; ns indicates p > 0,05, * indicates p < 0.05, ** indicates p < 0.01, *** indicates p < 0.001.

**Figure S2, related to Figure 2.**
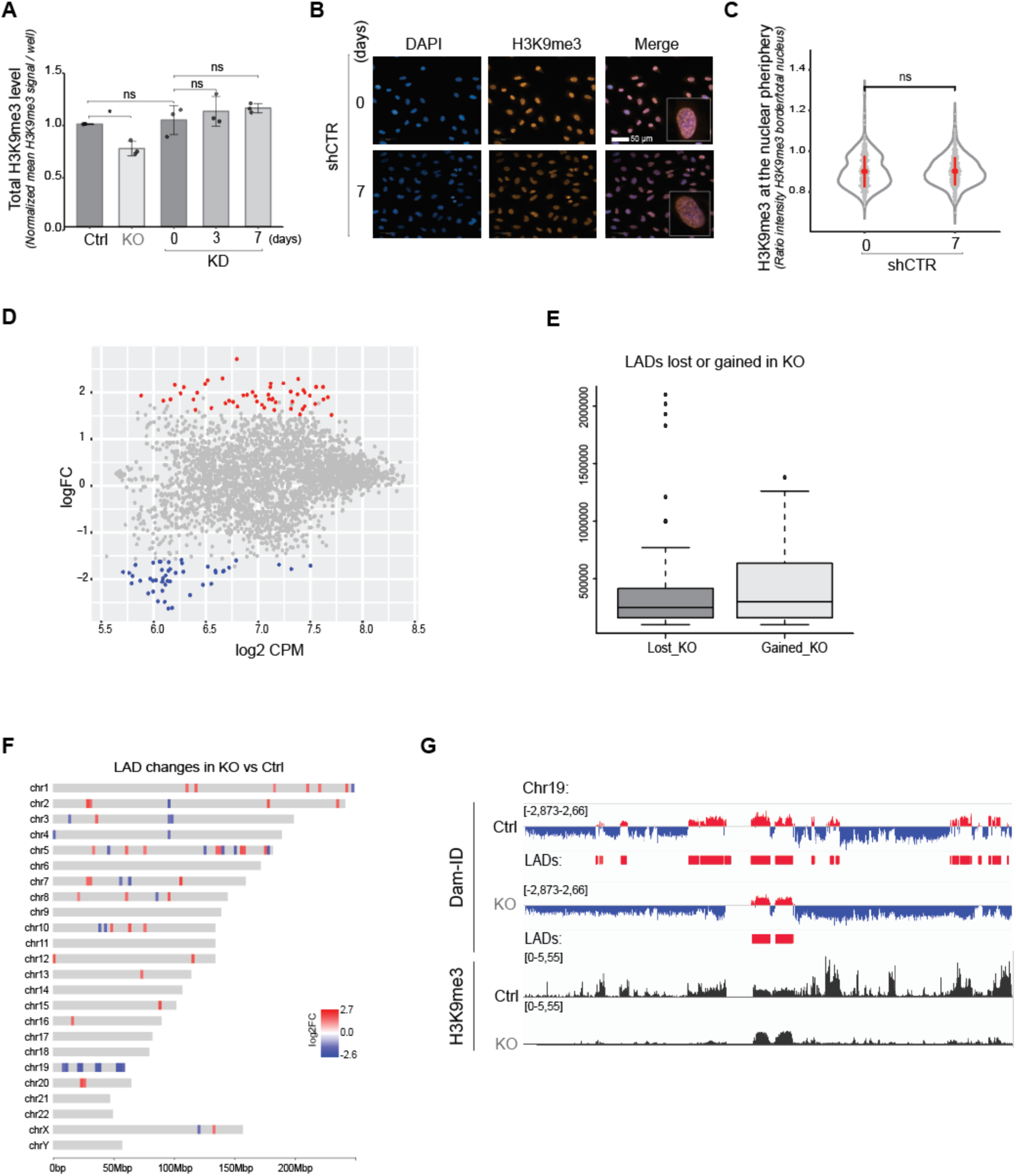
SETDB1 regulate the H3K9me3 distribution but does not impact the LADs composition. **A.** Quantification of the normalized to the Ctrl level of H3K9me3 in the nucleus in different cell lines normalized to Ctrl. Bars represent the mean of N=3 biological replicates +/− SD error bars, with individual values shown. Statistical significance was determined using an unpaired t-test; ns indicates p > 0,05, * indicates p < 0.05, ** indicates p < 0.01, *** indicates p < 0.001. **B.** Representative immunofluorescence images taken with the automated Operetta system used for the quantification for shCTR control cells and dox-treated cells. Cells stained with anti-H3K9me3 antibody, represented in orange. DNA was labelled with DAPI in blue. Scale bar: 50 µm. **C.** Quantification of H3K9me3 signal at nuclear periphery measured with the automated microscope (Operetta) corresponding to the phenotype observed in panel B. Each point represents the ratio between the intensity of the H3K9me3 signal at the nuclear periphery (defined by DAPI) and the total nuclear H3K9me3 signal intensity. 500 cells were randomly picked over more than 10 000 cells per condition. The red dot indicates the mean of N=3 biological replicates +/− SD error bars. Statistical significance was determined using the Wilcoxon test; ns indicates p > 0,05, * indicates p < 0.05, ** indicates p < 0.01, *** indicates p < 0.001, **** indicates p < 0.0001. **D.** MA plot depicting lamina-associated domains (LADs) ≥100 kb identified in Ctrl and SETDB1 KO cells. Each dot represents an individual LAD. Red and blue dots indicate LADs significantly gained or lost in SETDB1 KO relative to Ctrl cells (log_₂_ fold change > 1.5; false Discovery Rate, FDR < 0.05). **E.** Boxplot comparing the comparable genomic span of gained and lost LADs upon SETDB1 KO. Both categories span approximately 22 Mb. **F.** Graphical representation of the chromosomal distribution and positioning of gained (red) and lost (blue) Lamina-Associated Domains (LADs) in SETDB1 KO compared to Ctrl cells. The magnitude of change is indicated by a color gradient reflecting the log_₂_ fold change (log2FC). **G.** Integrated Genome Viewer (IGV) tracks displaying DamID-seq and ChIP-seq data from Ctrl and SETDB1 KO cells. Tracks represent Dam-Lamin B1 (DamID) and H3K9me3 (ChIP_H3K9me3) normalized signals, respectively. LADs are highlighted in red, while inter-LAD regions are shown in blue. The figure illustrates a loss of LADs on chromosome 19 (Chr19) in SETDB1 KO cells relative to Control, accompanied by a corresponding decrease in H3K9me3 signal. H3K9me3 ChIP-seq data were obtained from reference(Zakharova et al., 2022).

**Figure S3, related to Figure 3.**
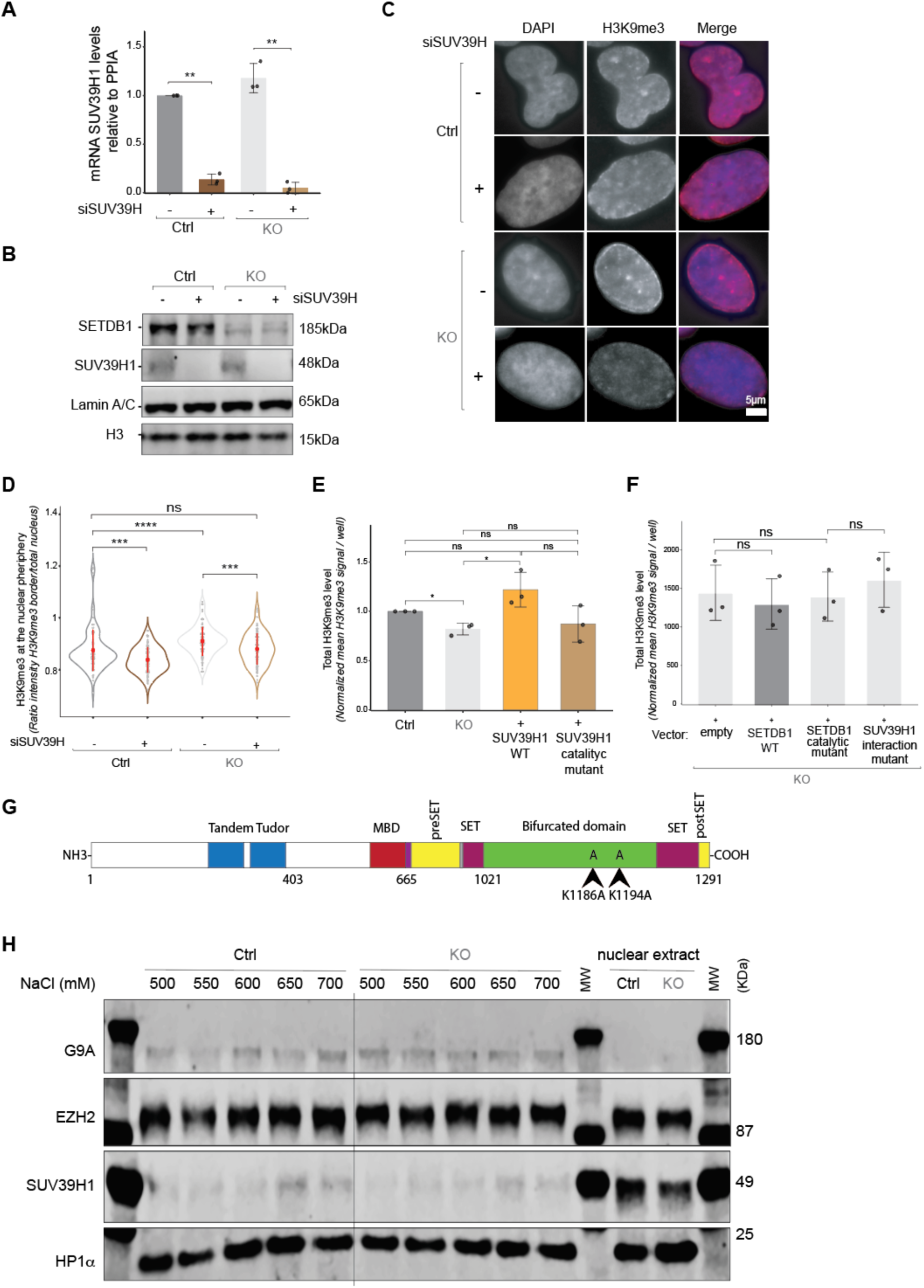
The SETDB1-SUV39H1 balance controls H3K9me3 distribution at the nuclear periphery. **A.** mRNA levels of SUV39H1 measured by RT-qPCR and normalized to PPIA. Expression levels are relative to A549 cells (Ctrl). siRNA against SUV39H is indicated as “+” and siCTR as “-”. Bars represent the mean of N=3 biological replicates +/− SD error bars, with individual values shown. Statistical significance was determined using an unpaired t-test; ns indicates p > 0,05, * indicates p < 0.05, ** indicates p < 0.01, *** indicates p < 0.001. **B.** Western blot analysis of SETDB1 and SUV39H1 protein levels in Ctrl and SETDB1 KO +/−siRNA against SUV39H cell lines. Lamin A/C and H3 were used as a loading control. **C.** Representative immunofluorescence images of Ctrl and SETDB1 KO +/− siRNA against SUV39H. Cells were stained with anti-H3K9me3 antibody. The nuclei were labelled with DAPI. The merged images show DAPI in blue and H3K9me3 in red. Scale bar: 5µm. **D.** Violin plot showing the quantification of H3K9me3 signal at nuclear periphery measured with the automated microscope (Operetta) corresponding to the phenotype observed in panel C. Each point represents the ratio between the intensity of the signal at the nuclear periphery (defined by DAPI) and the total nuclear H3K9me3 signal intensity. 100 cells were randomly picked over more than 10 000 cells per condition. The red dot indicates the mean of N=3 biological replicates +/− SD error bars. Statistical significance was determined using the Wilcoxon test; ns indicates p > 0,05, * indicates p < 0.05, ** indicates p < 0.01, *** indicates p < 0.001, **** indicates p < 0.0001. **E.** Bar plot showing the quantification of the total level of H3K9me3 in the nucleus in different cell lines normalized to Ctrl measured with the automated microscope (Operetta) corresponding to the phenotype observe*d* in panel C. Bars represent the normalized to the Ctrl mean of N=3 biological replicates +/− SD error bars, with individual values shown. Statistical significance was determined using an unpaired t-test; ns indicates p > 0,05, * indicates p < 0.05, ** indicates p < 0.01, *** indicates p < 0.001. **F.** Bar plot showing the quantification of the total level of H3K9me3 in the nucleus in different cell lines normalized to Ctrl measured with the automated microscope (Operetta) corresponding to the phenotype observe*d* in panel Fig. 3C. Bars represent the normalized to the Ctrl mean of N=3 biological replicates +/− SD error bars, with individual values shown. Statistical significance was determined using an unpaired t-test; ns indicates p > 0,05, * indicates p < 0.05, ** indicates p < 0.01, *** indicates p < 0.001. **G.** Structure of SETDB1 showing the position of the two lysins (K1186 and K1194) that were modified with alanine (A) to disrupt the SETDB1-SUV39H1 interaction. **H.** Solubility test for SUV39H1 in Ctrl and SETDB1 KO cells using increasing NaCl concentration (500 to 700 mM). G9A, EZH2 and HP1α are used as control to show that there is no change in solubility for these proteins between Ctrl and SETDB1 KO. On the right the total nuclei extract shows that the level of SUV39H1 is similar in Ctrl and SETDB1 KO cells.

**Figure S4, related to Figure 4.**
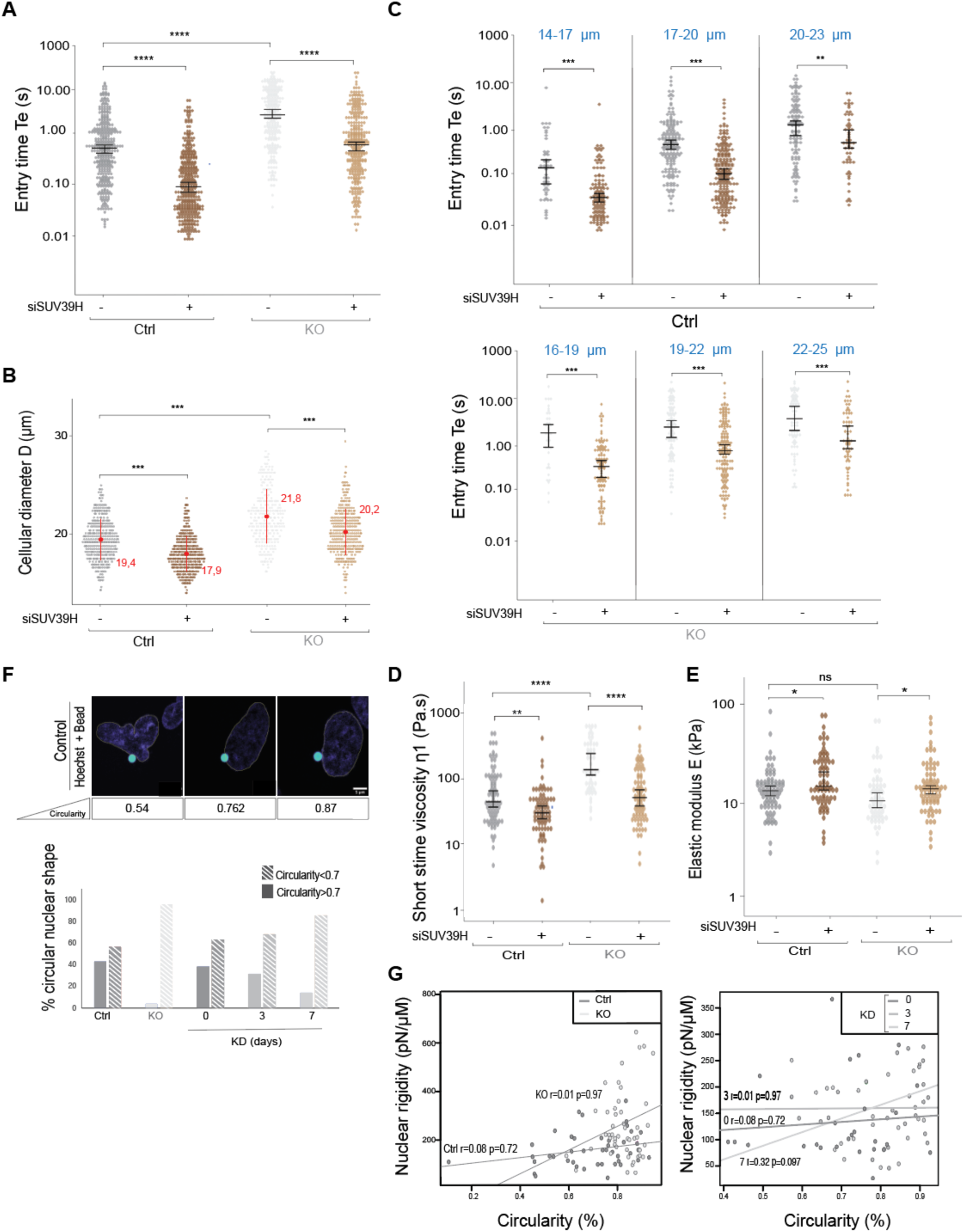
Morphological parameters alone cannot account for the observed biophysical changes. **A.** Dot plot representation of the entry time Te (in seconds, s) of Ctrl and SETDB1 KO cells treated with siCTR and siSUV39H. The dot plots of the non-Gaussian distributions are displayed on a logarithmic scale. Each dot represents a single cell; medians and 95% CI are shown. Data represent N = 3 biological replicates, each with more than 100 cells (n > 100). Statistical significance was determined using the Wilcoxon test; ns indicates p > 0,05, * indicates p < 0.05, ** indicates p < 0.01, *** indicates p < 0.001, **** indicates p < 0.0001. **B.** Dot plot representation of the cellular projected diameter D (as a proxy of the cell size, in μm) in Ctrl and SETDB1 KO cells treated with siCTR and siSUV39H. Each dot represents a single cell; mean +/−SD is shown in red. Data represent N = 3 biological replicates, each with more than 100 cells (n > 100). Statistical significance was determined using the Wilcoxon test; ns indicates p > 0,05, * indicates p < 0.05, ** indicates p < 0.01, *** indicates p < 0.001, **** indicates p < 0.0001. **C.** Distributions of entry time Te (in seconds, s) for Ctrl (top) and SETDB1 KO (bottom) cells treated with siCTR and siSUV39H sorted by cell diameter D in three groups. Each dot represents a single cell; medians and 95% CI are shown. Data represent N = 3 biological replicates. Statistical significance was determined using the Wilcoxon test; ns indicates p > 0,05, * indicates p < 0.05, ** indicates p < 0.01, *** indicates p < 0.001, **** indicates p < 0.0001. **D.** Dot plot representation of the short-time viscosity h _1_, in Pa.s, of Ctrl cells and SETDB1 KO cells treated with siCTR and siSUV39H. The dot plots of the non-Gaussian distributions are displayed on a logarithmic scale. Each dot represents a single cell; medians and 95% CI are shown. Data represent N = 3 biological replicates, each with more than 50 cells (n > 50). Statistical significance was determined using the Wilcoxon test; ns indicates p > 0,05, * indicates p < 0.05, ** indicates p < 0.01, *** indicates p < 0.001, **** indicates p < 0.0001. **E.** Dot plot representation of the elastic modulus E in kPa of Ctrl cells and SETDB1 KO cells treated with siCTR and siSUV39H. The dot plots of the non-Gaussian distributions are displayed on a logarithmic scale. Each dot represents a single cell; medians and 95% CI are shown. Data represent N = 3 biological replicates, each with more than 50 cells (n > 50). Statistical significance was determined using the Wilcoxon test; ns indicates p > 0,05, * indicates p < 0.05, ** indicates p < 0.01, *** indicates p < 0.001, **** indicates p < 0.0001. **F.** Circularity calculated with the Fiji program from confocal images taken during optical tweezers indentation experiments in Ctrl cells. Percentage of circular nuclei (circularity > 0.7) and non-circular nuclei (circularity<0.7) in Ctrl, SETDB1 KO, SETDB1 KD 0, 3d, 7d dox treated cells. **G.** Scatter plot showing correlation between nucleus rigidity and nucleus circularity in Ctrl and SETDB1 KO cells (left), and in the SETDB1 KD 0, 3d, 7d doxycycline-treated cells (right).

